# Different translation dynamics of β- and γ-actin regulates cell migration

**DOI:** 10.1101/2021.01.05.425467

**Authors:** Pavan Vedula, Satoshi Kurosaka, Brittany MacTaggart, Qin Ni, Garegin A. Papoian, Yi Jiang, Dawei Dong, Anna Kashina

**Affiliations:** Department of Biomedical Sciences, School of Veterinary Medicine, University of Pennsylvania, Philadelphia, PA; Institute of Advanced Technology, Kindai University, Kainan, Wakayama, Japan; Department of Chemical and Biomolecular Engineering, University of Maryland, College Park, MD; Department of Chemistry, University of Maryland, College Park, MD; Department of Mathematics and Statistics, Georgia State University, Atlanta, GA; Institute for Biomedical Informatics, Perelman School of Medicine, University of Pennsylvania,, Philadelphia, PA

## Abstract

β- and γ-cytoplasmic actins are ubiquitously expressed in every cell type and are nearly identical at the amino acid level but play vastly different roles *in vivo*. Their essential roles in embryogenesis and cell migration critically depend on the nucleotide sequences of their genes, rather than their amino acid sequence. However it is unclear which gene elements underlie this effect. Here we address the specific role of the coding sequence in β- and γ-cytoplasmic actins’ intracellular functions, using stable cell lines with exogenously expressed actin isoforms and their “codon-switched” variants. When targeted to the cell periphery using the β-actin 3′UTR, β-actin and γ-actin have differential effects on cell migration. These effects directly depend on the coding sequence. Single molecule measurements of actin isoform translation, combined with fluorescence recovery after photobleaching, demonstrate a pronounced difference in β- and γ-actins’ translation elongation rates, leading to changes in their dynamics at focal adhesions, impairments in actin bundle formation, and reduced cell anchoring to the substrate during migration. Our results demonstrate that coding sequence-mediated differences in actin translation play a key role in cell migration.

## Introduction

Actin is one of the most essential and abundant eukaryotic proteins, highly conserved across the tree of life. Among the six mammalian actins, β- and γ-cytoplasmic actins are the only two that are ubiquitously expressed in every vertebrate cell type and share the highest identity at the amino acid level, with only four conservative substitutions within their N-termini (Vandekerckhove and Weber, 1978). Despite this near-identity, the coding sequences for these two actin isoforms are different by approximately 13% due to synonymous substitutions (Erba et al., 1986). Our work has previously shown that this coding sequence difference can lead to differential arginylation of these two actins. Arginylated β-actin accumulates *in vivo*, while arginylated γ-actin is degraded (Zhang et al., 2010).

In mice, β- and γ-cytoplasmic actin play vastly different physiological roles. β-actin knockout leads to defects in embryogenesis and early embryonic lethality (Bunnell et al., 2011; Shawlot et al., 1998; Shmerling et al., 2005; Strathdee et al., 2008; Tondeleir et al., 2013; Tondeleir et al., 2014), while γ-cytoplasmic actin knockout mice survive until birth and have much milder overall phenotypic defects (Belyantseva et al., 2009; Bunnell and Ervasti, 2010). Our prior work showed that β-actin’s nucleotide sequence, rather than its amino acid sequence, underlies its essential role in embryogenesis (Vedula et al., 2017). Using CRISPR/Cas9, we edited the five nucleotides at the beginning of the β-actin coding sequence within the β-actin gene *(Actb)*, causing it to encode γ- actin protein (*Actbc-g*, β-coded-γ-actin). Such *Actbc-g* mice developed normally and showed no gross phenotypic defects, thus demonstrating that the intact β-actin gene, rather than protein, defines its essential role in organism’s survival (Patrinostro et al., 2018; Vedula et al., 2017). Thus, nucleotide sequence constitutes a major, previously unknown determinant of actin function, even though it is unclear which specific nucleotide-based elements of the actin gene play a role in this effect.

Here we tested whether the coding sequence alone, in the context of invariant non-coding elements, and independent of the positional effects of the actin gene, plays a role in cytoplasmic actins’ intracellular function. To do this, we incorporated the coding sequences of β- and γ- actin, and their “codon-switched” variants (β-coded-γ-actin and γ-coded-β-actin) into otherwise identical constructs containing the β-actin promoter, an N-terminal eGFP fusion, and the β-actin 3′UTR. Stable expression of these constructs in mouse embryonic fibroblasts (MEFs) resulted in dramatically different effects on directional cell migration. While cells expressing β-actin migrated at rates similar to wild type untransfected cells in wound healing assays, cells expressing γ-actin migrated nearly 2-fold faster. This difference was directly coding sequence-dependent, as evident by the use of the “codon-switched” actin variants. Expression of γ- and γ-coded-β-actin led to changes in cell morphology and distribution of the focal adhesions, which were larger in size and localized mostly at the cell periphery rather than under the entire cell. This phenotype has been previously linked to poorer cell attachment and faster migration (Kim and Wirtz, 2013). Focal adhesions in γ-actin and γ-coded-β-actin-expressing cells were often not visibly associated with long actin bundles. In contrast, long actin cables could be clearly seen anchoring focal adhesions in β-actin and β-coded-γ-actin-expressing cells.

Single molecule measurements of actin translation using the SunTag system at the focal adhesion sites showed a ~2-fold faster translation elongation of β- compared to γ-actin. Fluorescence recovery after photobleaching (FRAP) demonstrated that γ-actin accumulation in cells was slower than β-actin, further confirming global differences in actin isoform translation. Molecular simulations of actin assembly at the focal adhesions showed that differences in translation rates can directly impact actin bundle formation, leading to shorter actin bundles in the case of slower translating γ-actin, in agreement with our experimental data.

Our results demonstrate that, nucleotide coding sequence-dependent translation rates, coupled to zipcode-targeted actin mRNA localization, play an essential role in differential actin isoforms’ function in cell migration.

## Results

### β- and γ-actin coding sequences have differential effects on cell migration speed

To test the specific effect of coding sequence on intracellular functions of actin isoforms, we generated immortalized MEFs stably expressing β- and γ-actin coding sequences, as well as their codon-switched variants (β-coded-γ-actin and γ-coded-β-actin), cloned into identical expression constructs under the β-actin promoter, containing an N-terminal eGFP fusion and the β-actin 3′UTR (Fig. 1, top left). This construct design enabled us to confine our experiments to the effects of the coding sequence and exclude any potential contribution from other elements known to mediate differences between β- and γ-actin, including promoter-mediated transcription (Tunnacliffe et al., 2018), differential 3′UTR mediated mRNA targeting (Hill and Gunning, 1993; Katz et al., 2012), and differential N-terminal processing (Zhang et al., 2010). Cell populations stably expressing eGFP constructs were checked to ensure similar levels of eGFP mRNA, as well as to confirm that the expression of the exogenous eGFP-actin did not have any significant effect on the endogenous β- and γ-actin mRNA levels (Fig. S1). We also confirmed that β-actin 3′UTR targeted the eGFP-actin mRNA to the cell periphery, using fluorescence *in situ* hybridization (FISH) (Fig. S2). Finally, we confirmed that the level and distribution of F-actin in each of the cell lines was largely similar to each other (Fig S3). Thus, in these cell lines, the effects of exogenously expressed actin could be tested without perturbation of other actin-related processes that are essential for cell viability.

**Fig. 1.**
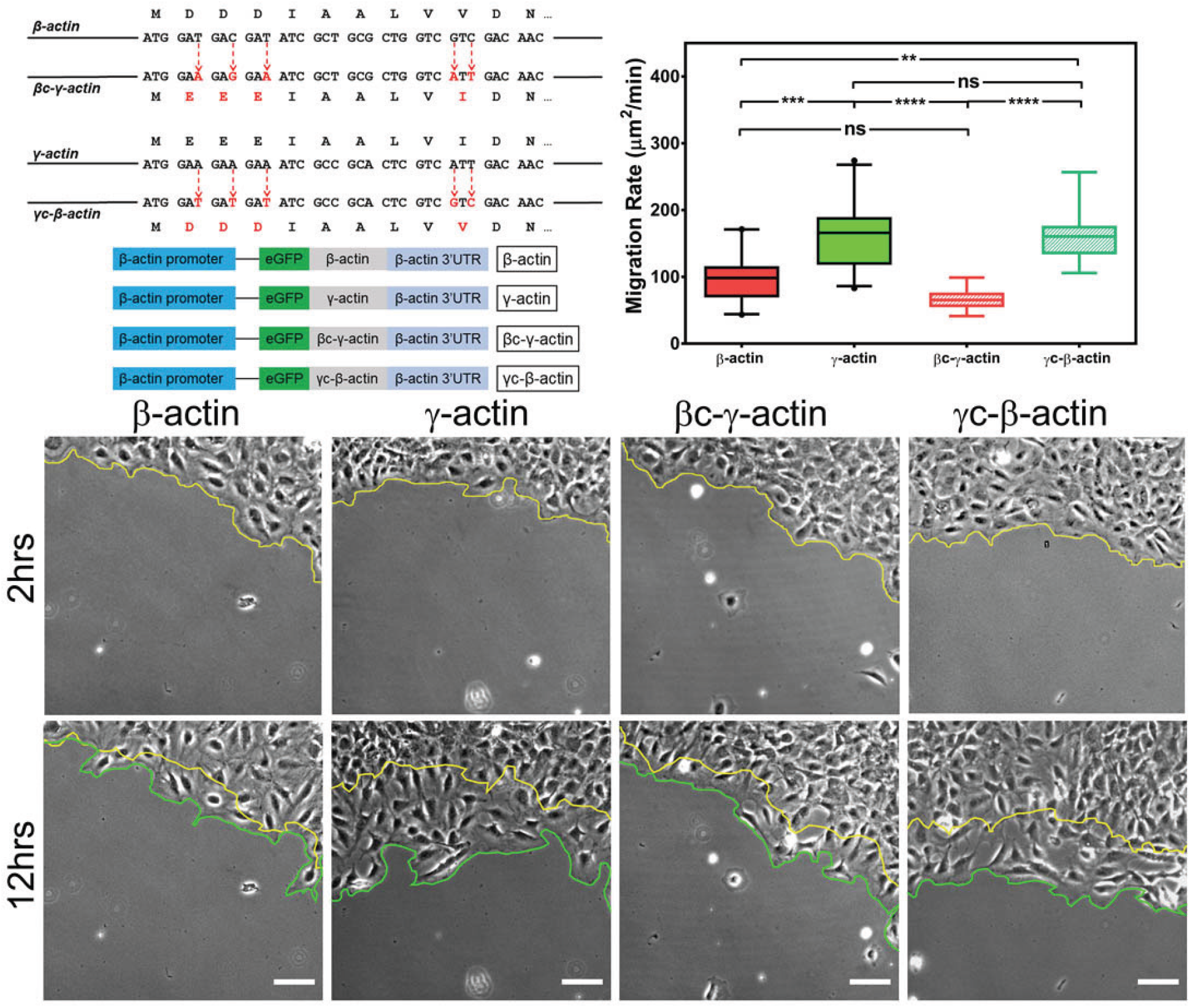
Cell migration speed is regulated by actin isoform coding sequence. Top left, mutagenesis strategy used to generate “codon-switched” actin variants and linear maps of the major constructs used in this study. Top right, box and whisker plots of migration rates of cell lines expressing different actin constructs listed on the top. Δ3′UTR denotes the constructs lacking of the β-actin 3′UTR responsible for zipcode targeting. All other constructs contained the β-actin 3′UTR. Bottom, representative images of the migrating wound edge, with the initial and the final position of the edge denoted by the yellow and the green line, respectively. Cell migration rates were derived as area covered over time by the entire cell monolayer (calculated as the area between the yellow and the green lines). N=20 (for β-actin); 22 (for γ-actin); 18 (for βc-γ-actin); 19 (for γc-β-actin). One way non-parametric ANOVA yielded a p value <0.0001 with multiple comparisons shown on the plot. Error bars represent SEM. *p<0.05, **p<0.01, ***p<0.001, and ****p<0.0001. See also supplementary videos S1-4. Scale bars, 100 μm.

β- and γ-cytoplasmic actin make up more than 50% of the total actin in these cells and have been previously shown to play major non-overlapping roles in directional cell migration (Patrinostro et al., 2017). We therefore tested whether cells expressing eGFP-β-actin or eGFP-γ-actin showed any differences in cell migration, using a wound healing assay. Strikingly, while cells expressing eGFP-β-actin migrated at rates similar to wild type untransfected cells, cells expressing eGFP-γ-actin migrated nearly 2-fold faster (Fig 1, top right and bottom; Fig. S4; Supplemental Videos 1 and 2). This difference in cell migration rates was coding sequence dependent, as seen in cells expressing the codon-switched actin variants, γ-coded-β-actin (which migrated faster, similarly to those expressing γ-actin) and β-coded-γ-actin (which migrated slower, like β-actin-expressing cells) (Fig. 1, top left, Supplemental Videos 3 and 4). Thus, the effect of actin isoform expression on directional cell migration is mediated by their nucleotide coding sequence and does not appear to be influenced by their amino acid sequence.

In normal cells, the β-actin 3′UTR contains a zipcode sequence which is required for its mRNA localization to the cell periphery and has been shown to be important for directional cell migration (Condeelis and Singer, 2005; Katz et al., 2012). γ-actin mRNA has no such sequence and does not undergo targeting to the cell periphery (Hill and Gunning, 1993). All our constructs described above contained the β-actin 3′UTR with the zipcode sequence as one of the constant elements. To test whether 3′UTR-mediated targeting of actin mRNA affects the cell migration phenotypes observed in our stable cell lines, we performed the same experiment using cell lines stably expressing similar actin constructs, but without the β-actin 3′UTR (Fig. S2). These cells did not exhibit significant differences in cell migration rates (Fig. S5). Thus, differences in the effects of cytoplasmic actin coding sequences on cell migration require mRNA targeting to the cell periphery.

### Actin isoforms affect focal adhesion size, cell spreading, and actin dynamics at the focal adhesions in a coding sequence-dependent manner

Changes in cell migration rates are normally associated with changes in actin dynamics at the leading edge, rate and persistence of leading edge protrusions and retractions, as well as focal adhesion formation and dynamics, which affect cell spreading, polarization, and attachment to the substrate. Focal adhesions’ strength and persistence is closely regulated by their association with actin filaments, which grow at the focal adhesion sites to form a dynamic actin bundle that participates in anchoring the cells to the substrate. Thus, focal adhesions critically depend on actin dynamics in the vicinity of the adhesion site. In turn, focal adhesions can regulate cell spreading and polarization, in addition to cell migration rates.

To test these processes in eGFP-actin isoform-transfected cells, we first looked at the rate and persistence of leading edge protrusions and retractions, but found no consistent differences between the cell lines that correlated with cell migration rates (Fig. S6). We next assessed focal adhesion dynamics in these cells using Total Interference Reflection Fluorescence Microscopy (TIRF-M) of eGFP-β-actin and eGFP-γ-actin. Since imaging volume in TIRF-M is limited to the basal 300 nm or less, we reasoned that most of the actin signal visible in this volume should be associated with focal adhesion patches. Imaging the long-term (hours) behavior of actin at focal adhesion patches during wound healing using TIRF-M revealed that in migrating cells, eGFP-β-actin patches appeared more prominent and persisted considerably longer than eGFP-γ-actin patches (Fig. 2A, B), suggesting that focal adhesions in eGFP-β-actin-expressing cells persist for longer periods of time. At the same time, testing short-term (5 min) actin dynamics at the focal adhesions using fluorescence recovery after photobleaching (FRAP) showed no notable differences in focal adhesion recovery rates that correlated with either coding or amino acid sequence (Fig S7). Thus, different actin isoforms affect long term focal adhesion persistence without strongly affecting short term focal adhesion or protrusion dynamics during persistent directional migration at the cell leading edge.

**Fig. 2.**
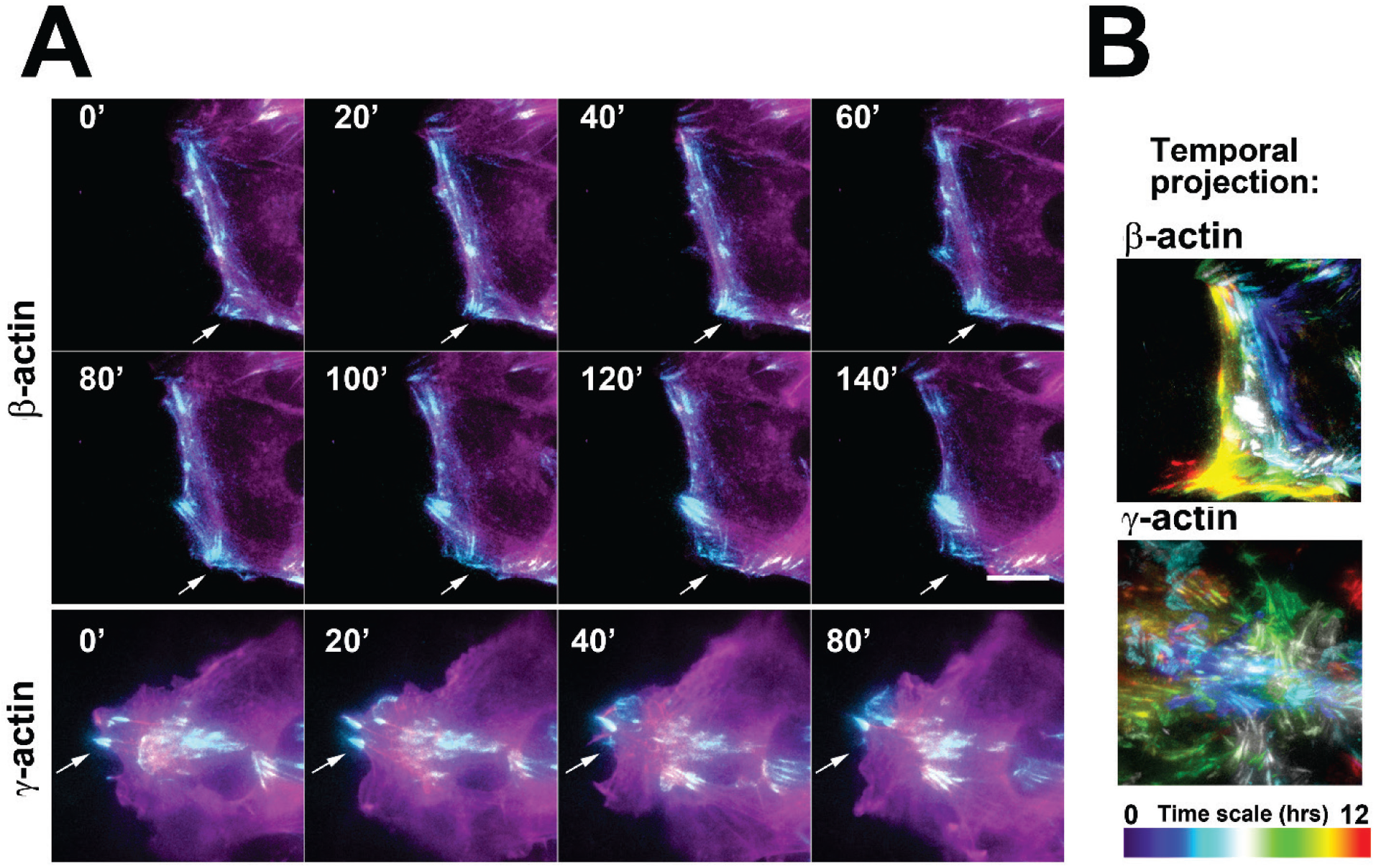
Actin isoforms have differential effect on focal adhesion dynamics. A, Montages of eGFP-β-actin (top two rows) and eGFP-γ-actin (bottom row) expressing cells undergoing wound healing following a scratch wound. TIRF images (Cyan) are overlaid with widefield images (Magenta). Scale bar = 10μm. Arrows point to focal adhesions being formed and disassembled over time. B, Maximum intensity projections of TIRF-M images of eGFP-actin over time for each of the cell lines during a 12-hour wound healing. Scale represents the temporal color scale. Note that β-actin persists longer in the TIRF plane as compared to γ-actin.

To get deeper insights into the focal adhesion morphology and distribution in cells transfected with different actin isoforms, we used single cell cultures, in which cells are grown at low density, and thus the morphology and cytoskeleton-dependent structures in individual cells are much easier to visualize. Notably, cells in such scarce cultures are under no stimuli to migrate, and thus many of them remain stationary or move randomly around the same area, resulting in much slower rates of persistent migration and overall displacement over time. Consequently, such sparsely grown cells transfected with different actin isoforms do not prominently differ from each other in their migration (Fig. S8), even though they are expected to undergo similar actin isoform-related changes at the subcellular level.

To analyze focal adhesions and spreading in these cells, we first used TIRF-M to image single cells stained with antibodies to the focal adhesion protein paxillin. These assays revealed prominent differences in focal adhesion morphology and distribution between the cell lines (Fig. 3, top row of images; see also figures S9-S12). In eGFP-β-actin expressing cells, focal adhesions had normal elongated morphology and were distributed throughout the entire cell footprint. In contrast, eGFP-γ-actin expressing cells formed focal adhesions that appeared more rounded and localized mostly at the cell periphery. This trend depended on the actin coding sequence: focal adhesions in eGFP-β-coded-γ-actin expressing cells resembled those in eGFP-β-actin, while focal adhesions in eGFP-γ-coded-β-actin expressing cells were similar to those in eGFP-γ-actin expressing cells. In addition, imaging eGFP-actin in widefield showed that most focal adhesions in cells expressing eGFP-β-actin and eGFP-β-coded-γ-actin were associated with long thick bundles of actin emanating from the focal adhesion point (Fig. 3, top, and figures S9-S12), while such dorsal bundles connecting to the focal adhesions were much less prominent in γ-actin and γ-coded-β-actin expressing cells.

**Fig. 3.**
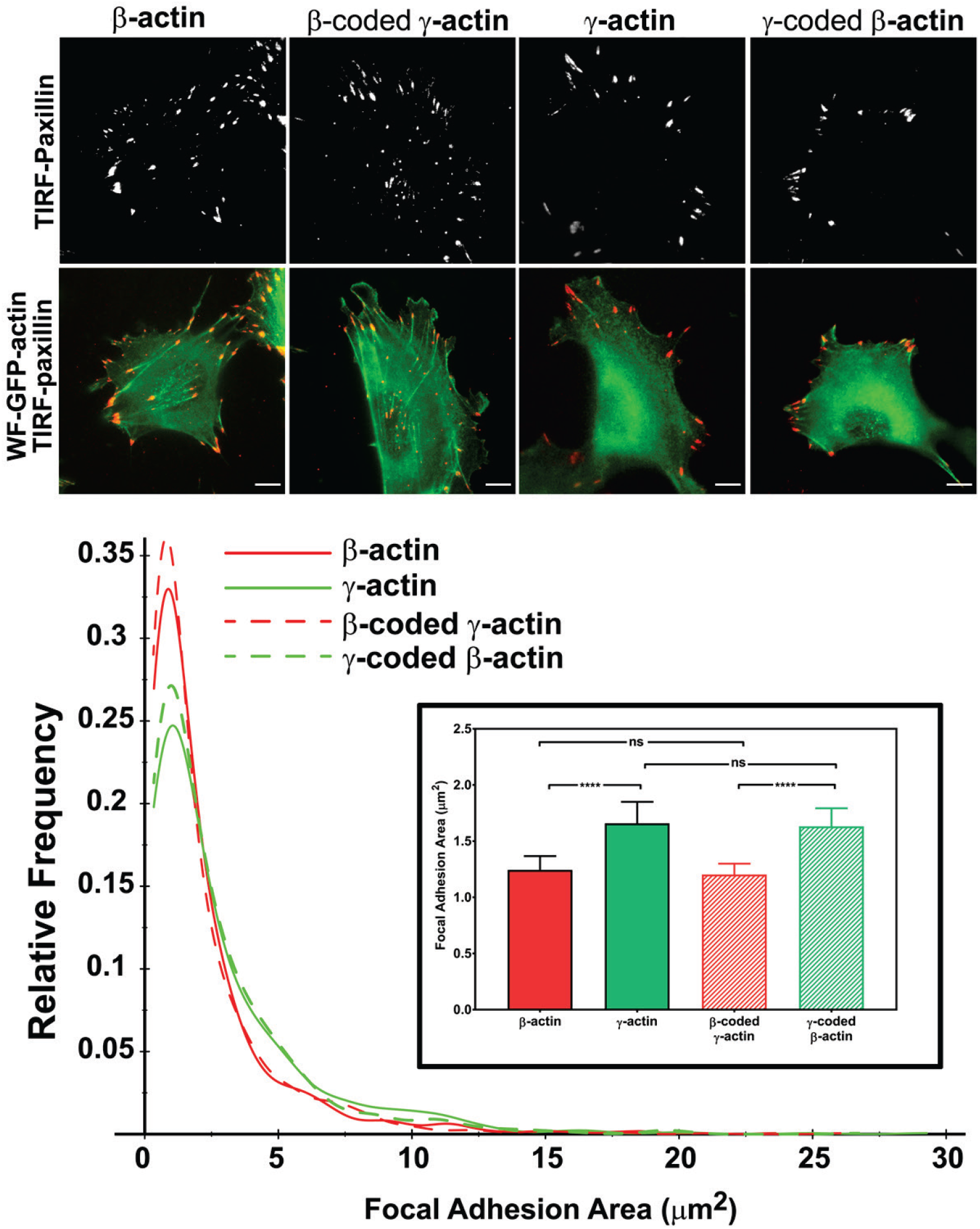
Actin isoform coding sequence affects the overall size and distribution of focal adhesions. Top, representative images of eGFP-actin transfected cells stained with anti-paxillin to visualize focal adhesions. TIRF mages of paxillin staining are shown alone (top) and as overlay with wide field eGFP signal (bottom). Bottom, quantification of focal adhesion size in the four cell lines, shown as a distribution plot and as a bar chart in the inset. Scale bar = 10μm. N = 996, 786, 1245, and 1041 focal adhesions for the respective cell types from right to left from ≥ 20 cells for each were used for the analysis. Results for one-way ANOVA with multiple comparison are indicated on the graph.

Focal adhesion size uniquely predicts cell migration rate (Kim and Wirtz, 2013), with the larger focal adhesions correlating with faster migration speeds. We measured focal adhesion area in all four cell lines, and found that faster migrating cells expressing γ-actin, and γ-coded-β-actin, indeed had significantly larger focal adhesions than slower migrating cells expressing β-actin, and β-coded-γ-actin (Fig. 3, bottom). Thus, β- and γ-actin coding sequences determine the size and distribution of focal adhesions in migrating cells in a manner that correlates with changes in their migration speed.

We also measured focal adhesion recovery rates in single cell cultures using FRAP. Focal adhesions in cells transfected with β-actin and β-coded-γ-actin recovered slightly faster than cells transfected with γ-actin and γ-coded-β-actin (Fig. S13). While statistically significant, these coding sequence-dependent differences appear small, and thus it is unclear if they can prominently contribute to the cells’ phenotype.

Cell spreading and polarization are critically determined by their adhesion to the substrate and correlate with their migratory behavior (Kim and Wirtz, 2013). To test whether focal adhesion changes in our cell lines are accompanied by changes in cell spreading and polarization, we used Celltool (Pincus and Theriot, 2007) to analyze shape distribution of single cells stably expressing eGFP-β-actin, eGFP-γ-actin, and their codon-switched variants. Using images of live single cells, we parameterized the shape space into various shape modes, and selected two modes that accounted for ~60% of variance in shapes across all cells (Fig. S14, inset). The first mode roughly captures variation in the size of the cell footprint on the substrate and accounts for ~40% of variance in shape, while the second mode captures cell polarization and accounts for ~20% of the variance in shape. Using these shape modes to analyze images of cells expressing different actin isoforms’ coding sequences, we found that cells expressing eGFP-β-actin had a larger footprint (Fig. S14, clustered to the left of the y-axis) and had more variance in their polarization (Fig. S14A, spread across the y axis), while cells expressing eGFP-γ-actin exhibited the opposite trends (S14A, clustered mostly in the top right quadrant). Expression of the codon-switched actin variants, β-coded-γ-actin, and γ-coded-β-actin, showed that the footprint size depended on the actin isoform coding sequence, while the polarization variance appeared to be amino acid sequence-dependent. Changes in the area of the cell footprint can arise due to either reduced cell spreading or reduced overall cell size. To distinguish between these possibilities, we quantified the area of trypsinized near-spherical cells (pre-spreading), which directly reflects cell size and volume. Cells expressing the γ-actin coding sequence were slightly smaller than those expressing β-actin coding sequence (Fig. S14B, left). This ~6% difference in cell size was far less prominent than the difference in spread cell area (Fig. S14B, right), which accounted for a greater than 80% change in the size of cell footprint. Thus, cells expressing γ-actin are less spread on the substrate, and this difference in spreading is coding sequence-dependent.

### β-actin exhibits faster intracellular translation elongation than γ-actin

In search for an underlying mechanism that could link actin isoforms’ coding sequence to their intracellular properties, we turned to our previous study that used computational predictions of the mRNA secondary structure for β- and γ-actin. This study suggested that the coding region of β-actin mRNA forms a more relaxed secondary structure than that of γ-actin, predicting potential differences in translation elongation rate (Zhang et al., 2010). Such differences, if prominent enough, could in principle lead to changes in cells’ ability to form focal adhesions and migrate. To test this prediction, we first compared the rates of overall protein accumulation of eGFP-β- and eGFP-γ-actin, by comparing FRAP of the total eGFP signal in the cell after whole-cell photobleaching. We reasoned that this would serve as a proxy for estimation of newly synthesized β- and γ-actin (Fig 4A). Notably, the recovery observed in these FRAP experiments within a 10-minute imaging window arises from the folding and maturation of already synthesized eGFP fused to actin (since the eGFP maturation rate *in vivo* has been estimated to be approximately 14 minutes (Balleza et al., 2018; Iizuka et al., 2011)); given the constant time delay, this recovery rate directly reflects the rate of *de novo* synthesized actin accumulation within the imaging window. Photobleaching was calibrated to ensure cells remained healthy and visually normal during the experiment (Fig. 4A, bottom). The recovery rate was significantly faster for eGFP-β-actin compared to eGFP-γ-actin (Fig. 4A, top right). Thus, newly synthesized β-actin accumulates in cells faster than γ-actin.

**Fig. 4.**
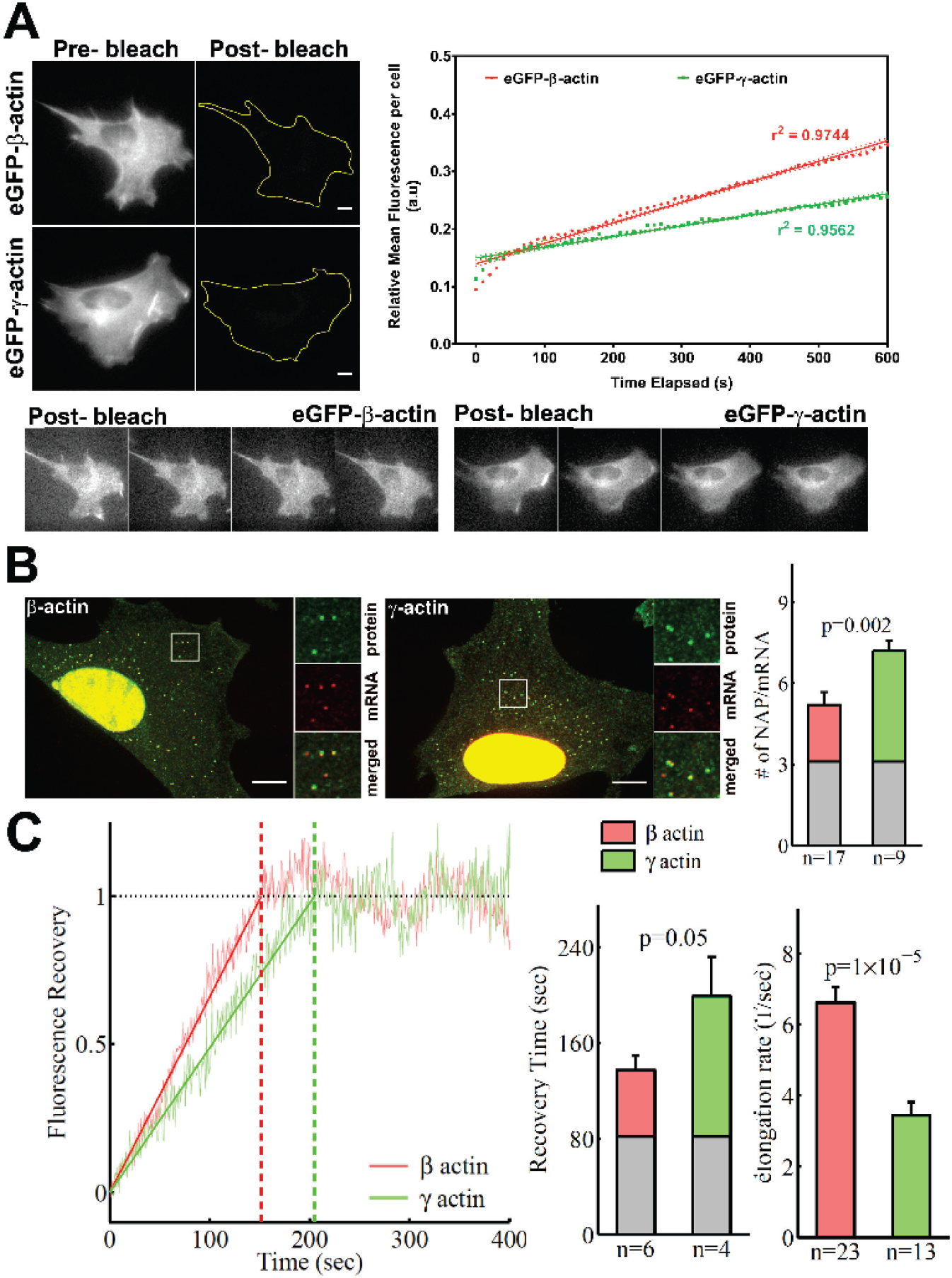
β-actin accumulates in cells faster than γ-actin and exhibits faster translation elongation. A. Images and quantification of GFP fluorescence in cells before and after photobleaching the entire intracellular GFP signal. Top left panels show representative cells before and after photobleaching, with gray levels scaled to show the actual difference in signal intensity. Top right graph shows the fluorescence recovery after photobleaching (FRAP) curve during the first 10 minutes post-bleach. Data points are dots and the linear regression curves are in bold (with dotted lines representing the 95% confidence bands). N = 8 for eGFP-β-actk and N = 10 for eGFP-γ-actin. Bottom, post-bleached images of cells taken at 2.5′ intervals from 0′ (left) to 10′ (right), with the gray levels scaled up to enhance the residual eGFP signal. Scale bars = 10 μm. B. Left: representative images of actin protein and mRNA used for the experiments. Insets on the right of each image show the enlarged region indicated in the image on the left. Right: Quantification of ribosomes per mRNA, calculated as the SunTag signal from the nascent peptides (NAP) at each translation site divided by the HaloTag signal from the mRNA. C. Left, FRAP curves showing recovery of the green fluorescence signal in individual translation spots. Right, fluorescence recovery time for each actin isoforms and translation elongation rate (amino acids/s) calculated from the fluorescence recovery and NAP/RNA. In B right and C right: gray boxed area at the bottom of each bar indicates the contribution from SunTag, AID and linker portion of the construct (see Methods), and p-values are from one-tail Welch’s t-test. Error bars represent SEM for n data points, calculated geometrically and plotted in linear fashion.

To directly estimate β- and γ-actin translation elongation rates, we performed single molecule imaging of nascent peptide synthesis (SINAPS) for these two actin isoforms using the SunTag system (Wu et al., 2016). Similarly to the constructs used for eGFP-actin stable cell line generation, we ensured that the coding sequence was the only variable, flanked by otherwise identical upstream and downstream elements, including the UbC promoter for constitutive expression, the N-terminal 5′ SunTag fusion to visualize the nascent peptide, the C-terminal auxin-induced degron to degrade fully synthesized polypeptides and reduce the background signal, the β-actin 3′UTR for cell periphery targeting, and MS2 repeats in the non-coding region to visualize mRNA via constitutively expressed MS2 coat binding protein (MCP) fused to a HaloTag (Fig. S15).

Simultaneous imaging of SunTag-bound superfolder GFP (sfGFP) and MS2-bound HaloTag in fixed cells (labelled with JaneliaFluor 646) enabled us to estimate the number of nascent peptides (NAPs) per mRNA, a measure of the ribosome load and, by proxy, translation elongation rate. Assuming similar translation initiation rates for both constructs (predicted based on their identical upstream and downstream elements and N-terminal fusions), fewer elongating ribosomes in this assay arise from their faster translocation over the mRNA, leading to weaker sfGFP signal per mRNA. Thus, differences in ribosome load per mRNA (and thus, the number of NAPs per mRNA) would directly indicate differences in translation elongation.

γ-actin coding sequence showed nearly 2-fold higher level of NAPs/mRNA than that of β-actin (Fig. 4B, S16), indicating that the elongating ribosomes had 2-fold higher load, and thus slower translocation rate, over γ-actin mRNA compared to β-actin. This is the first characterization of different ribosome loads on individual mRNAs of two protein isoforms that have the same coding sequence length.

Next, we estimated the real-time translation elongation rate of β- and γ-actin using FRAP of individual translation sites. For this experiment, to minimize mRNA movement, we tethered SINAPS-β- and SINAPS-γ-actin mRNA to focal adhesions, by co-transfecting cells with vinculin-MCP-HaloTag fusion. Fluorescence recovery rate of individual translation sites in this experiment directly reflects the rate of translation elongation to generate new nascent peptides bearing the new sfGFP bound to the SunTag peptides. This recovery rate was ~2-fold slower for γ-actin compared to β-actin (Fig. 4C), confirming that γ-actin translation elongation is indeed slower than that of β-actin, in agreement with the NAP/mRNA measurements.

Using both sets of data, we conclude that the translation elongation rate of the two actin isoforms differs by approximately 2-fold – faster for β-actin compared to γ-actin (Fig. 4D).

### Faster initial subunit supply rates at the focal adhesions facilitate longer actin bundle formation

During cell migration, the initial formation of nascent focal adhesions critically depends on local actin subunit supply rate. Many studies assume that this subunit supply rate is not a limiting factor *in vivo*, due to high concentrations of G-actin at the cell leading edge (Raz-Ben Aroush et al., 2017), however increasing evidence suggests that not all of this actin is locally polymerization competent. It is possible that at a given moment some, or most, of the free actin can be sequestered, e.g., by monomer binding proteins (Skruber et al., 2018), forcing the elongating leading edge filaments to depend on de novo synthesized actin. In support, actin mRNA targeting to the cell leading edge is essential for cell migration, suggesting that local actin synthesis at the cell leading edge must be important (Katz et al., 2012). Furthermore, local actin translation bursts have been observed in neurons (Buxbaum et al., 2014). It is possible that these bursts, regardless of the overall actin concentrations, are required for locally supplying actin subunits at the focal adhesions during cell migration. If so, replacing the faster translationally elongating β-actin with the slower elongating γ-actin at these sites could potentially limit this supply and make a difference in focal adhesion anchoring, leading to smaller actin bundles at the focal adhesions, poorer spreading, and faster migration seen in γ-actin expressing cells.

Since measurements of local polymerization-competent actin in a cell are impossible to perform experimentally, we used the computational model of active networks, MEDYAN (Popov et al., 2016), to simulate actin bundle growth at the focal adhesions at different subunit supply rates, in the presence of non-muscle myosin II motors and α-actinin as crosslinkers, which are critical for actin filaments bundling in cells (Chandrasekaran et al., 2019) (Fig.5A and S17). In these simulations, filaments elongate by incorporating newly supplied actin monomers, and then bundle together through the action of myosin motors and crosslinkers. During 10-minute simulations – the time window typically sufficient for establishment of robust focal adhesions -- varying actin subunit supply rate resulted in pronounced differences in the length of the actin bundle growing from the focal adhesion site (Supplemental Videos 5 and 6). A 2-fold decrease in subunit supply rate resulted in over a 2-fold decrease in actin bundle length (Fig. 5A, right and Fig. S17B). To test this prediction experimentally, we measured the length of eGFP-actin decorated bundles emanating from paxillin-positive focal adhesion patches in cells stably expressing different eGFP-actin isoforms, using both the migrating cells at the edge of the wound and single cells (Fig. 5B, left). In both types of cultures, actin bundles associated with the focal adhesion sites were markedly longer in β-actin-expressing cells, compared to those expressing γ-actin (Fig. 5B, right). Moreover, these trends followed the actin coding sequence, rather than amino acid sequence (Fig. 5B). Thus, slower subunit supply dictated by differences in translation elongation rates of β- and γ-actin coding sequences during the initial events of focal adhesion formation and maturation bears direct consequences to cell adhesion and migration.

**Fig. 5.**
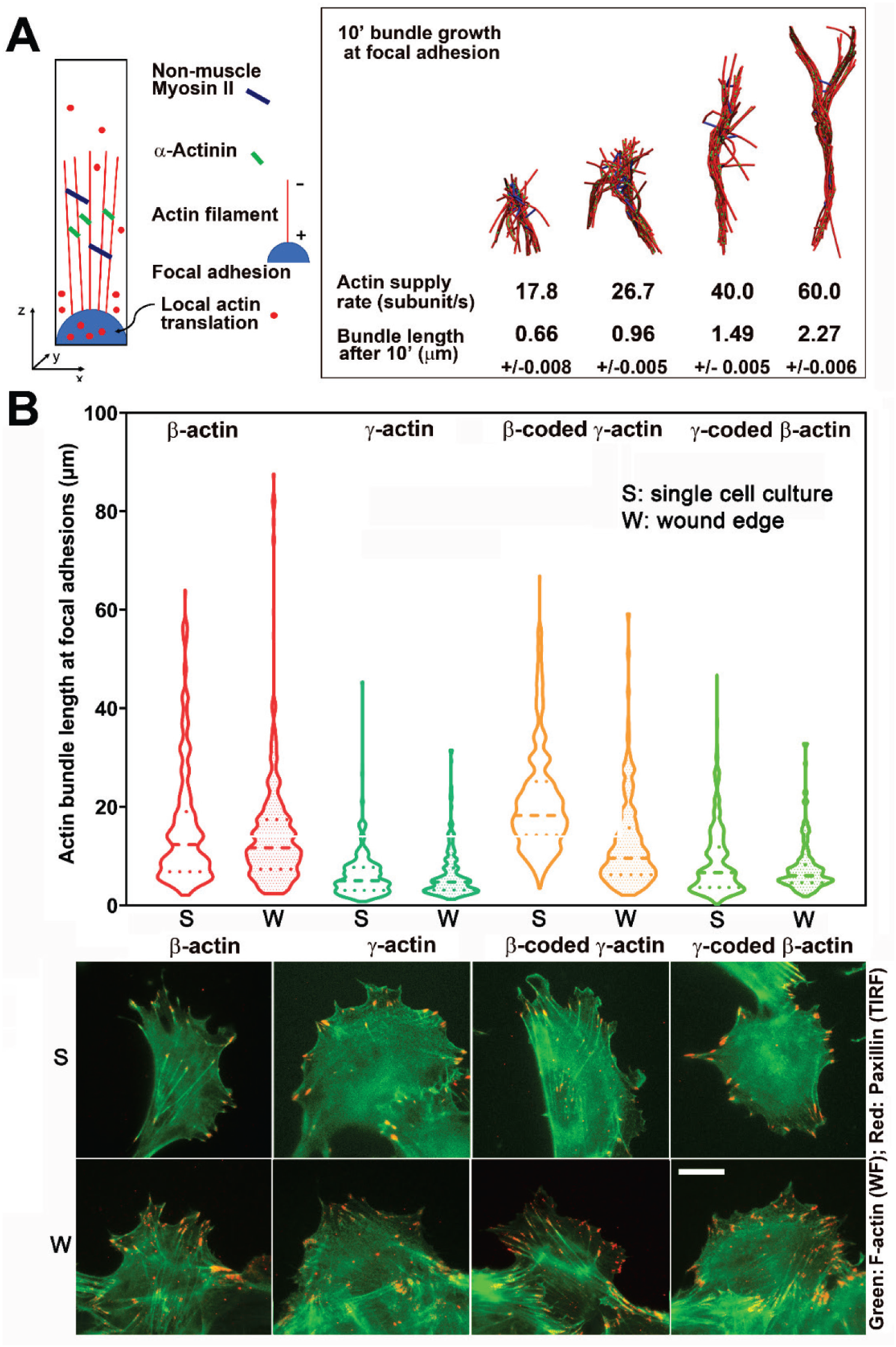
Faster β-actin subunit initial supply rate facilitates actin bundle formation at the focal adhesions to support normal cell migration. A. Molecular simulations of actin filament growth at the focal adhesion at four different subunit supply rates, using the components listed on the right. Length of the actin bundle for each supply rate is indicated underneath, with SD listed for five independent simulations. All systems contain 0.012 μM non-muscle myosin II mini-filaments and 1.25 μM alpha-actinin crosslinkers. B. (Top), Violin plots indicating the distribution of lengths of actin bundles emanating from focal adhesion. Horizontal dashed lines indicate median ± 95% CI. Actin bundle lengths measured from both single cell sparse cultures (S) and the cells at the migrating wound edge (W) during a wound healing assay are shown. Number of actin cables measured: (S) – β-actin = 1715, γ-actin = 756, β-coded-γ-actin = 482, and γ-coded-β-actin = 2383; (W) – β-actin = 1165, γ-actin = 1129, β-coded-γ-actin = 1726, and γ-coded-β-actin = 898. (Bottom) Representative images of the single cell cultures (S) and cells at wound edge (W) showing eGFP-actin (green) co-stained with focal adhesion marker paxillin (red). Scale bar, 20 μm.

## Discussion

Our study follows up on the recent discovery of the essential role of nucleotide, rather than amino acid, sequence in non-muscle actin isoform function and demonstrates for the first time that actin coding sequence, uncoupled from other gene elements, can directly affect cell behavior. We find that differences in β- and γ-actin coding sequences result in different ribosome elongation rates during their translation, leading to changes in cell spreading, focal adhesion anchoring, and cell migration speed. This study constitutes the first direct comparison of translation rates of two closely related proteins and the first demonstration that these translation rates can mediate their functions *in vivo*.

On the surface, it appears to be surprising that expression of the slower-translating γ-actin can make the cells move faster than the faster-translating β-actin. However, this result fits well into the context of the previously proposed localized bursts of β-actin translation, implicated in cell spreading and focal adhesion formation (Condeelis and Singer, 2005; Katz et al., 2012). Our data show that faster translating β-actin is required for generating stable focal adhesions, while the slower translating γ-actin leads to faster focal adhesion turnover. We propose that slower translation at the leading edge makes γ-actin less capable of supporting and sustaining strong focal adhesions, potentially by reducing the availability of actin for rapid bundle formation and anchoring of the cell-substrate attachment sites. In support, in γ-actin expressing cells, most focal adhesions, while larger in area, do not appear to be visibly anchored by prominent actin bundles. In contrast, cells expressing β-actin contain long actin cables emanating from most of the focal adhesion sites. This difference is highly likely to impair spreading and induce cells to glide over the substrate rather than migrating in a normal step-by-step manner characteristic of this cell type. We propose that this gliding, caused by weaker cell-substrate attachment, is responsible for the faster migration speeds in cells expressing γ-actin (Fig. 6). Notably, this increase in cell speed is only evident in a wound healing assay, where cells are collectively stimulated to migrate directionally, rather than randomly move around as typical for single cell cultures of MEFs. It is likely that actin behavior in response to the strong signals for cells to polarize and move directionally during wound healing depend on local actin translation more critically than during random migration, where cells can change directions or remain stationary for extended periods of time and are not constrained in the direction of their polarization and motility. These constraints could potentially involve cell-cell adhesions and other forms of signaling in dense cultures.

**Fig. 6.**
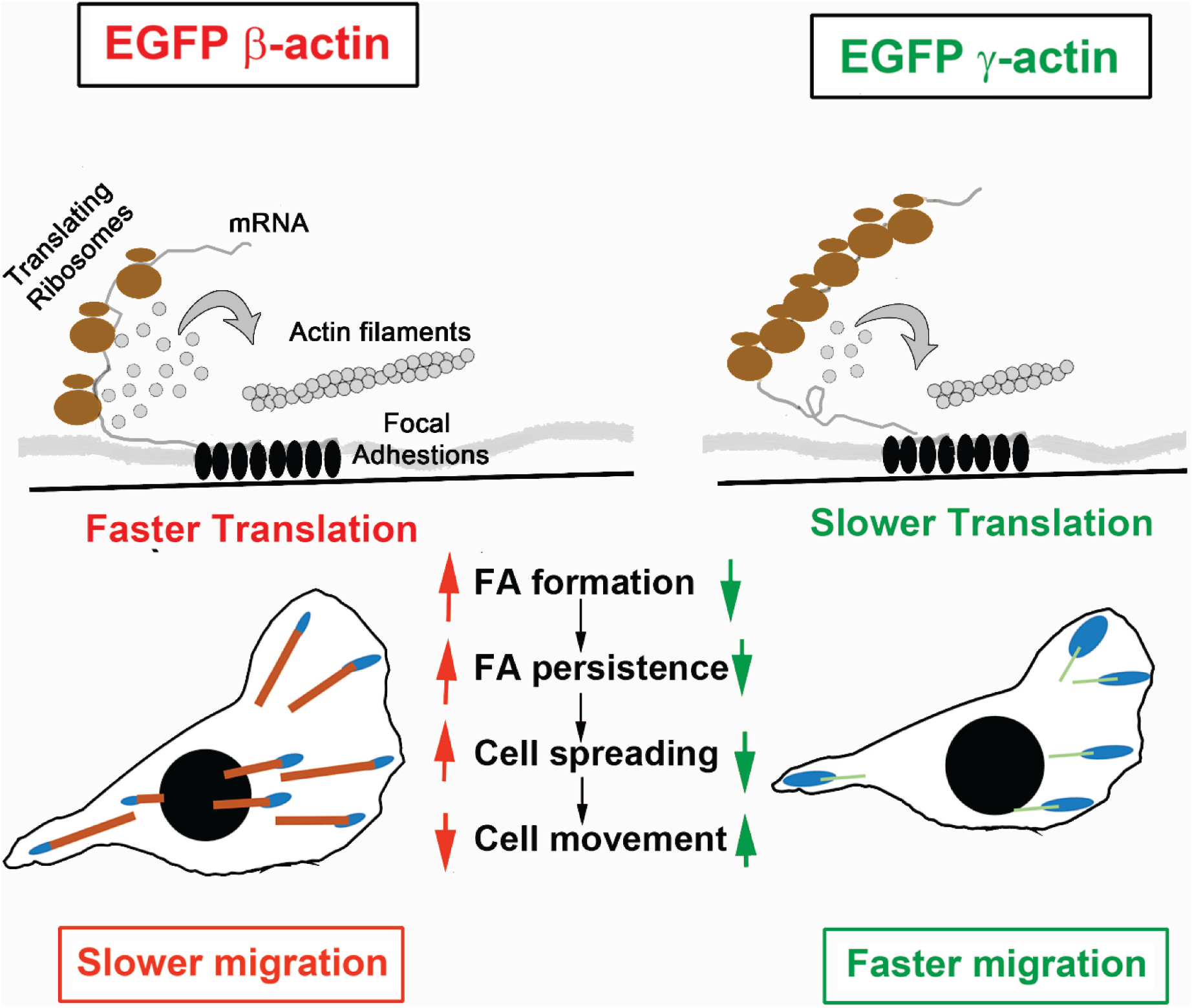
Hypothesis of cell migration regulation by actin isoforms’ translation rates. Faster translating β-actin facilitates focal adhesion (FA) formation and persistence, resulting in increased cell spreading and reduced rate of directional cell movement. Slower translating γ-actin has the opposite effect.

While the local supply rate of actin subunits to the forming focal adhesion sites during cell migration is nearly impossible to measure experimentally with the currently available methods, the use of modeling and simulation enables us to vary this parameter and estimate the subunit supply rate that could make a difference in this process. It appears surprising that a 2-fold difference in translation elongation rate found in our study could exert such a pronounced effect on the length of the actin bundles forming locally at the focal adhesion sites, especially given the fact that polymerization-competent actin should exist everywhere in the lamellipodia. Our results suggest that the concentration of this polymerization-competent actin pool may be far lower than previously estimated, potentially due to the action of monomer sequestering proteins or posttranslational modifications that may prevent actin from incorporating into filaments (Skruber et al., 2018). It is also possible that, even with higher actin concentration in the lamellipodia, its local diffusion rate cannot fully compensate for the differences in local translation between β- and γ-actin. Notably, these differences are expected to be even higher with the native non-eGFP-fused actin isoforms, which likely differ in their translation initiation rates in addition to the difference in elongation rates we observed. These questions, and the exact interplay between newly synthesized and diffusible actin in cells, constitute an exciting direction of future studies.

Our previous work showed that actin coding sequence leads to differential arginylation of β- and γ-actin (Zhang et al., 2010), and this arginylation is important for directional cell migration (Karakozova et al., 2006). In the present study, the use of N-terminal eGFP fusions likely excludes arginylation as a variable in our cell lines, since arginylation is believed to require an exposed β-actin N-terminus. The use of GFP-actin fusions also likely limits the types of effects we can observe, since these fusions are not fully functionally equivalent to the native actin and are impaired, e.g., in formin nucleation (Chen et al., 2012). Thus, our constructs cannot substitute for all aspects of normal actin in cells, and this could explain the fact that the strongest effects we observe are related to cell adhesion and migration, the processes that are likely able to fully utilize eGFP-actin. Notably, in our cell lines the endogenous β- and γ-actin are still present and still able to support cell migration, likely compensating for these types of functions, and potentially diminishing the observed phenotypes.

Thinking of the nucleotide sequence, rather than the amino acid sequence, as a determinant of actin function is a novel view that opens up many exciting questions. The present study demonstrates that coding sequence alone can play a significant role in cell behavior, but it does not exclude the possibility that other nucleotide-based elements of the actin gene also contribute to actin isoforms’ global role in organism survival. For example, it has been shown that a unique intron in the γ-actin gene can contribute to its function (Lloyd and Gunning, 1993). Studies of the interplay of nucleotide- and amino acid-based determinants in actin functions constitute exciting future directions in the field.

## Materials and Methods

### Constructs

Constructs were generated using the eTC GFP beta-actin full length plasmid, which was a gift from Robert Singer (Addgene plasmid # 27123) (Rodriguez et al., 2006). The human β-actin promoter and the TC-eGFP on the N-terminus were preserved. Mouse full length β-actin cDNA with the 3′UTR (eGFP-β-actin) or the coding sequence without 3′UTR (eGFP-β-actin Δ3′UTR) was cloned in place of human β-actin. β-actin coding sequence was replaced by γ-actin coding sequence in these two constructs to generate eGFP-γ-actin and eGFP-γ-actin Δ3′UTR. To make codon-switched constructs, point mutations were introduced and two additional constructs were generated eGFP-βcoded-γ-actin, and eGFP-γ-coded-β-actin.

pUbC-FLAG-24xSuntagV4-oxEBFP-AID-baUTR1-24xMS2V5-Wpre was a gift from Robert Singer (Addgene plasmid # 84561) and was used to generate the actin isoform SINAPS reporters (Wu et al., 2016). The β- and γ-actin coding sequences were cloned in place of the oxEBFP sequence in the original construct. For fixed cell imaging of ribosome load per mRNA, phage UbC NLS HA stdMCP stdHalo – a gift from Jeffrey Chao (Addgene plasmid # 104999) (Voigt et al., 2017), was used. For constructing the mRNA thether in live cell real time imaging of translation dynamics, pUbC-nls-ha-stdMCP-stdGFP – a gift from Robert Singer (Addgene plasmid # 98916) (Wu et al., 2015), was used to clone mouse vinculin sequence upstream of stdMCP, and stdGFP was replaced by HaloTag. pHR-scFv-GCN4-sfGFP-GB1-NLS-dWPRE – a gift from Ron Vale (Addgene plasmid # 60906) (Tanenbaum et al., 2014) was used for the NAP sensor. pBabe TIR1-9myc – a gift from Don Cleveland (Addgene plasmid # 47328) (Holland et al., 2012) was used for degrading the fully synthesized SINAPS construct.

### Generation of stable cell lines

Spontaneously immortalized Mouse Embryonic Fibroblasts (MEFs) were cultured in DMEM (Gibco) supplemented with 10% FBS (Gibco). Cells were transfected with the linearized EGFP-actin constructs described above. Following G418 selection, GFP positive cells were sorted using fluorescence activated cell sorting (FACS) and cultured. Live cell imaging was carried out in FluorBrite DMEM (Life Technologies) culture media supplemented with 10% FBS (Sigma), and L-glutamine (Gibco).

HEK-293T cells were cultured in DMEM (Gibco) supplemented with 10% FBS (Gibco). Lipofectamine 2000 (Life Technologies) was used to transfect these cells with plasmids for generating either lentiviral particles – pMD.G, pPAX, and plasmid containing gene of interest, or retroviral particles – pCL10A, and pBabe TIR1-9myc. Virus containing medium was harvested and used to infect immortalized MEFs in the following order: first, either UbC-NLS-HA-stdMCP-stdHalo for fixed cell imaging of number of NAPs/mRNA or Vinculin-stdMCP-Halo for live cell dynamics of translation elongation, second, TIR1-9myc followed by Puromycin selection of infected cells, third, scFv-GCN4-sfGFP-GB1-NLS, and lastly, either SINAPS-β-actin or SINAPS-γ-actin containing lentivirus. These polyclonal cells were used for imaging number of NAPs on each of β- and γ-actin constructs. Indole-3-acetic aicd was used at 500μg/mL to induce degradation of fully synthesized SINAPS constructs. Janelia Fluor 646 tagged Halo ligand (Promega) was used at 200nM final concentration to label SINAPS-mRNA in cells prior to fixation/imaging.

### Cell migration assays and imaging

Cell migration was stimulated by making an infinite scratch wound. The cells were allowed to recover for a period of 2 hours before imaging. Images were acquired using a 10X phase objective on a Lecia DMI 4000 equipped with a Hamamatsu ImagEM EMCCD camera. Images were captured every 5 minutes for 10 hours. Migration rates were measured as the area covered by the edge of the wound in the field of view per unit time using Fiji (NIH). For TIRF wound healing experiments, cells were imaged on a Nikon Ti with a 100X, 1.49 NA objective using the 488nm laser, and an Andor iXon Ultra 888 EMCCD.

### Fluorescence Recovery After Photobleaching

For all FRAP experiments, imaging was carried out on a Nikon Ti inverted microscope equipped with either an Andor iXon Ultra 888 EMCCD camera (0.13μm/pixel – for imaging TIRF-FRAP and Widefield whole cell eGFP-actin FRAP) or Andor iXon Ultra 897 EMCCD camera (0.16μm/pixel – for imaging SINAPS-FRAP using a Yokogowa CSU X1 spinning disc confocal). Photobleaching was carried out with a Bruker miniscanner equipped with XY Galvo mirrors. The region of interest for photobleaching was defined using a freehand ROI manager in NIS elements: an elliptical region encompassing an actin patch at the cell periphery for TIRF-FRAP, the entire cell for widefiled whole cell FRAP, and single pixel spot containing the translation site for SINAPS-FRAP.

eGFP-actin expressing cells were seeded on Matek glass bottom dishes and allowed to spread overnight. For photobleaching, the 488nm laser was set to 80% power and used to bleach a defined eGFP-actin patch at the cell periphery with a dwell time of 400μs/pixel. Images were acquired in the TIRF mode with the 488nm laser set to 50% power, and 200ms exposure and EM gain of 200. Images were acquired at 3s intervals for 12s pre-bleach and 6min post-bleach. Change in fluorescent intensity in a circle within an actin patch that was not bleached was used as a reference to account for photobleaching during acquisition. The change in intensity within a circle of the same area within the bleached actin patch was used to calculate the recovery curve. The obtained values were normalized to 1 at pre-bleach and the resulting post-bleach curves were fit using non-linear regression to a single exponential fit in GraphPad PRISM.

For whole cell eGFP-actin FRAP, the whole cell was outlined. For photobleaching, the 488nm laser was set to 70% power with a dwell time of 70μs/pixel. Acquisition was carried out using a 488nm LED illumination from Spectra/Aura with 10% illumination intensity, and 200ms exposure with an EM gain of 300. Images were acquired every 10 seconds for a total of 10 minutes after bleaching. The recovery curves obtained were fit using a linear regression model in GraphPad PRISM.

For live cell SINAPS-FRAP, cells expressing SINAPS-actin constructs thethered using Vinculin-stdMCP-Halo (See Plasmids and Generation of stable cell lines sections above). For photobleaching, the 488nm laser was set to 80% power with a dwell time of 1ms/pixel. Images were acquired in the spinning disc confocal mode with the 488nm laser set to 30% power, and 35ms exposure an EM gain of 300. Images were acquired at 700ms intervals for 30sec pre-bleach and 7min post-bleach.

### Translation elongation rate measurements

FishQuant (Mueller et al., 2013) was used to detect NAP and mRNA signals. Spots in the NAP channel that were within 300nm of a spot in the mRNA channel were considered bonafide NAP and used for estimating the integrated fluorescence intensity in both channels.

Airloclaize (Lionnet et al., 2011) was used to fit the signal from thethered NAPs. The integrated signal was recorded pre-bleach and post-bleach. These values were used to calculate the translation elongation rates of the the two actin isoforms.

Assuming that beta-actin has the same elongation rate as Suntag, AID and linkers, following the theoretical derivation of Wu et al. 2016, it is straight forward to show that the proportion of none beta-actin contribution to recovery time and NAP/mRNA is (L+(N+1)/2)/(S+L+(N+1)/2), in which N is the total number of Suntags, S is the beta-actin length in the unit of one Suntag, and L is the length of AID and linkers in the unit of one Suntag, shown as the gray bar in Figure 4B and 4C. It is not surprising to see from those figures that the ratios of recovery time to NAP/mRNA are similar for beta- and gamma-actin, since the initiation rates for both constructs should be the same given the identical N-terminal as well as C-terminal. Therefore, we calculated the variance weighted geometric average of the two ratios, which is T~26.9s, and used it to combine the data from recovery time and NAP/mRNA to calculate the beta-actin elongation rate: Rb=(S+L+(N+1)/2)/t or Rb=(S+L+(N+1)/2)/n/T, where t is recovery time and n is NAP/mRNA, for each data point, followed by geometric averaging. The contribution from Suntag, AID and linkers to recovery time is T0=(L+(N+1)/2)/Rb which is used to calculate the gamma-actin elongation rate: Rg=S/(t-T0) or Rg=S/(n*T-T0) for each data point, followed by geometric averaging. The results are shown in Figure 4D.

### Immunofluorescence staining and analysis

To quantify the amount of actin polymer, cells were seeded on coverslips in six well plate at 20,000 cells/well overnight and fixed in 4% (w/v) PFA at room temperature for 30 minutes. Cells were then stained with Phalloidin conjugated to AlexaFluor 594 (Molecular Probes). Images were acquired on Leica DM6000 at 40X and the total intensity of phalloidin per cell was measured using Fiji (NIH). To analyze focal adhesions in single cells, eGFP-actin-expressing cells were seeded on coverslips allowed to adhere and spread overnight. Cells were then fixed in 4% (w/v) PFA at room temperature for 30 minutes followed by 0.5% Triton-X 100 treatment for 5 minutes. Cells were incubated with mouse anti-paxillin monoclonal antibody (BD Biosciences), followed by AlexaFluor 555 conjugated goat-anti-mouse secondary antibody (Life Technologies). Cells were imaged with Citifluor (Cytoskeleton Inc.) anti-bleaching agent. To analyze cell spreading and cell area, Celltool was used to outline cell shapes and classify them and extract shape modes. The shape modes that captured 60% of the overall variability in the shape model were used to assess the distribution of cell shapes in a PCA plot. Additionally, a kernel density estimate of the marginal was used to plot the area of focal adhesions.

### Fluorescence *in situ* hybridization

eGFP mRNA probes (conjugated to Quasar 670 dye) were purchased from LGC Biosearch Technologies (VSMF 1015-5) and fluorescence *in situ* hybridization was carried out as per manufacturers’ protocol. Briefly, cells were seeded onto coverslips in six well plate at 20,000 cells/well overnight and fixed in 4% (w/v) PFA at room temperature for 30 minutes followed by treatment with 70% alcohol at 4°C for 1 hour. Cells were incubated with 125nM probes at 37°C overnight. Cells were stained with DAPI (5ng/mL) and mounted using Prolong Diamond (Life Technologies). Images were acquired using Leica DM6000 at 40X. Z-stacks were acquired, and blind deconvolution was carried out using Leica LAS X software.

### Real Time PCR

Cells were seeded onto 10cm culture dishes and grown to confluence. RNA was isolated using RNeasy mini kit (Qiagen) and cDNA was synthesized using oligo dT primers first strand cDNA synthesis kit (Applied Biosystems). After standard curves were obtained, qPCR was carried out using SybrGreen (Applied Biosystems) and the following primer sets. PCR was carried out on QuantStudio Flex 6 Real Time PCR system (Applied Biosystems). ΔΔCt method was used to estimate the relative expression levels of mRNA using TBP as the reference gene.

β-actin

Forward primer: 5′ GATCAAGATCATTGCTCCTCCTG 3′
Reverse primer: 5′ AGGGTGTAAAACGCAGCTCA 3′
γ-actin

Forward primer: 5′ GCGCAAGTACTCAGTCTGGAT 3′
Reverse primer: 5′ TGCCAGGGCAAATTGATACTTC 3′
eGFP

Forward primer: 5′ GTGAAGTTCGAGGGCGACA 3′
Reverse primer: 5′ TCGATGTTGTGGCGGATCTT 3′
TBP

Forward primer: 5′ TAATCCCAAGCGATTTGCTGC 3′
Reverse primer: 5′ AGAACTTAGCTGGGAAGCCC 3′

### Simulations of actin filament growth at the focal adhesions

Computational simulations of actin bundle growth from focal adhesions to predict the bundle length at different β- and γ-actin local supply rates were performed using a recently developed software MEDYAN (Popov et al., 2016). In brief, MEDYAN simulates actin networks by integrating the stochastic diffusion-reaction dynamics and mechanical relaxation of the cytoskeletal network. Diffusing molecular species, including actin monomers, unbound myosin motors and unbound crosslinkers, are contained in a solution phase. Stochastic chemical reactions such as actin (de)polymerization and (un)binding of motors and linkers follow mass-action kinetics, and change the mechanical energy of the actin network. The net forces are then periodically relaxed using conjugate gradient mechanical equilibration. This step also updates reaction rates of motor walking, motor unbinding, and linker unbinding, based on residue tension after minimization.

We used a 1×1×4 μm^3^ simulation box, containing non-muscle myosin II motors, alpha-actinin crosslinkers, actin monomers, and actin filaments growing from the bottom focal adhesion region. The focal adhesion region was presented as a hemisphere with 30 actin filaments attached. We tested that the actin filaments never grow longer than 4μm in the z-direction, and thus, no length constraints on the actin bundles factored into the simulations. Actin filaments were only allowed to elongate at one end (the barbed end), while the elongation rate constant was averaged over filament polymerization rates and depolymerization rates of both barbed end and pointed end. The filament elongation was driven by the addition of actin monomers to the system, simulating the synthesis of actin monomers near the focal adhesion region. Multiple actin supply rates were tested at 50% increments based on the experimental measurements of the differences between β- and γ-actin synthesis rate. The simulations were run for 10 minutes to match the timescale of the experiments. The starting concentration of actin at the attachment site was assumed to be ~2 μM locally, creating an initial bundle at around 0.1 μm long. In the simulation, the majority (more than 90%) of actin for the filament growth was assumed to arise from the de novo subunit addition. The concentrations of myosin mini-filaments (0.012 μM to 0.021 μM) and alpha-actinin crosslinkers (1.25μM) were chosen to ensure proper bundling of filaments (all model parameters are listed in Table S1).

To determine the actin bundle length, we measured the F-actin distribution along the Z-axis and defined the actin bundle length as the width of central 80% of the F-actin distribution (Fig. S17A). Although the length measured in simulations was much shorter than the experiments, the beta-actin bundles were ~50-80% longer than gamma-actin bundles, in agreement with the experimental measurements.

## Acknowledgements

We thank members of the Kashina lab for helpful and stimulating discussions. This work was supported by NIH grants R35GM118017 and R01NS102435 to A.K., R01CA201340 and R01EY028450 to Y.J. Simulations of actin bundle growth at the focal adhesion are a result of the collaboration between Kashina and Jiang labs initiated at the NSF-sponsored workshop funded by Award No. MCB-1411898. Simulations were supported by NSF grant CHE-1800418 and PHY-1806903 to G.P. and were performed on Deepthought2 Supercomputer at the University of Maryland.

