## Supplemental Online Information for "Different translation dynamics of β- and γ-actin regulates cell migration"

[1]Department of Biomedical Sciences, School of Veterinary Medicine, University of Pennsylvania, Philadelphia, PA; [2] Institute of Advanced Technology, Kindai University, Kainan, Wakayama, Japan; [3]Department of Chemical and Biomolecular Engineering, University of Maryland, College Park, MD; [4]Department of Chemistry, University of Maryland, College Park, MD; [5]Department of Mathematics and Statistics, Georgia State University, Atlanta, GA; [6]Institute for Biomedical Informatics, Perelman School of Medicine, University of Pennsylvania, , Philadelphia, PA

Vedula_et_al_Fig.S1

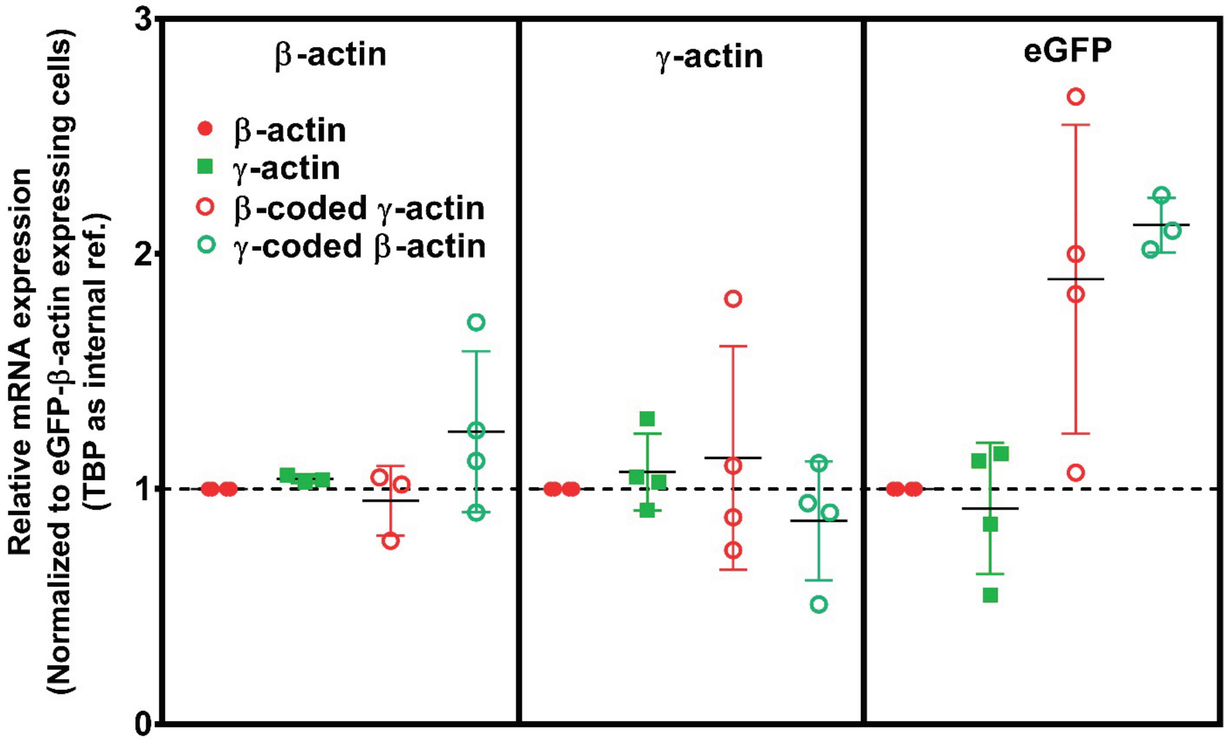


**Fig. S1, related to Fig.1. Cell transfection with actin isoforms does not perturb the endogenous levels of actin mRNA.** A, Real time PCR of the four cell lines, using primer pairs detecting actin isoforms' endogenous 3'UTRs and GFP. Each of the qPCR was repeated 4 times and outliers were excluded in Graphpad PRISM. The error bars represent 95% CI. One way ANOVA indicated there was a significant change only in the amount of eGFP mRNA ($p = 0.02$), with multiple comparison showing a significant change between β-actin and γ-coded β-actin only ($p = 0.02$).

**Vedula_et_al_Fig.S2**

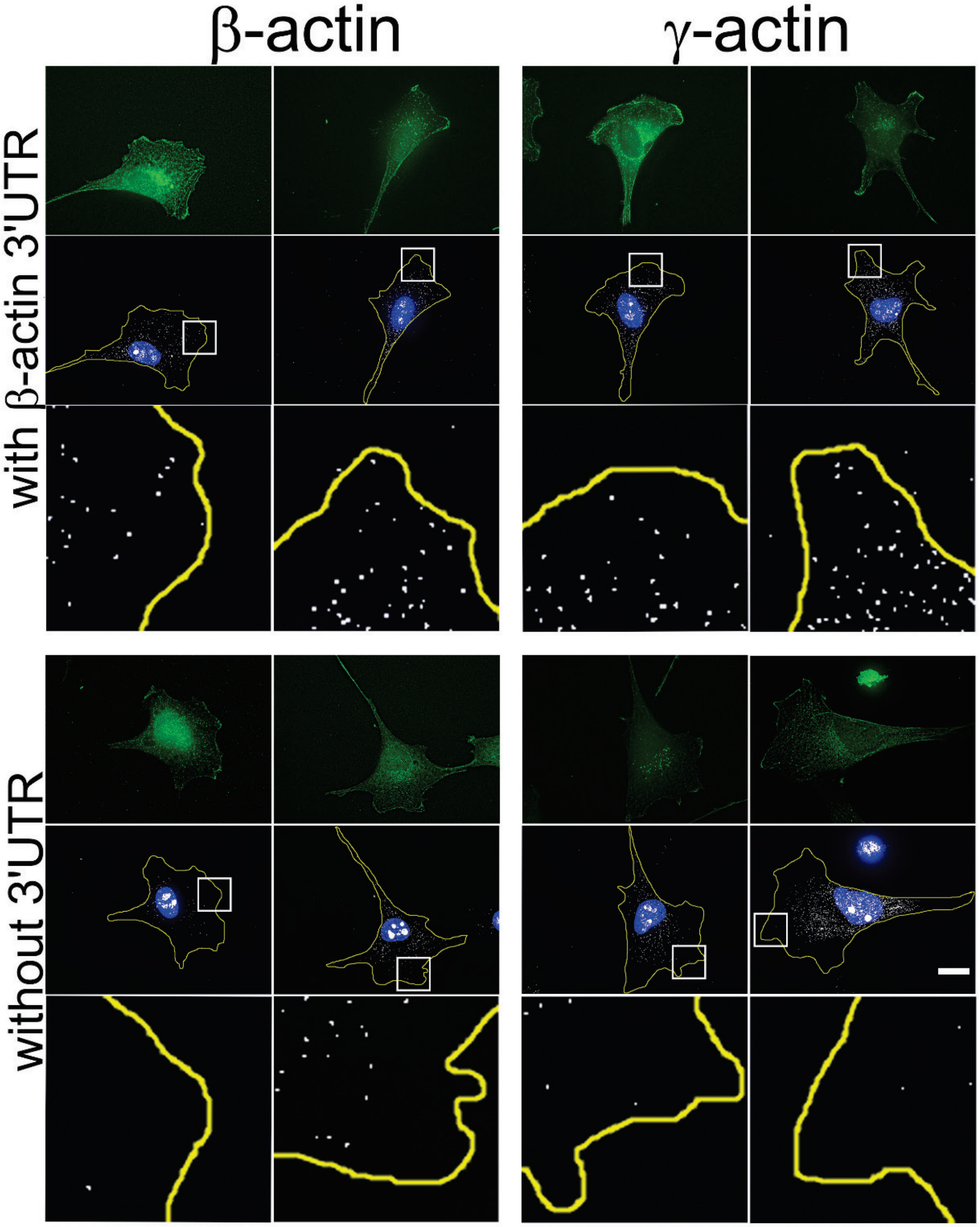


**Fig. S2, related to Fig. 1. Transfected actin isoform mRNAs are targeted to the cell periphery via 3'UTR.** Images of GFP-actin transfected cells lines with and without 3′UTR (as indicated on the left) visualized by GFP fluorescence (green), DAPI (blue), and GFP-encoding mRNA (white) detected using single molecule fluorescence in situ hybridization (smFISH) probes against GFP mRNA. Insets show a zoom of the boxed areas in FISH images, showing localization of eGFP mRNA significantly more pronounced in the presence of β-actin 3′UTR. Scale bar, 20 μm.

**Vedula_et_al_Fig.S3**

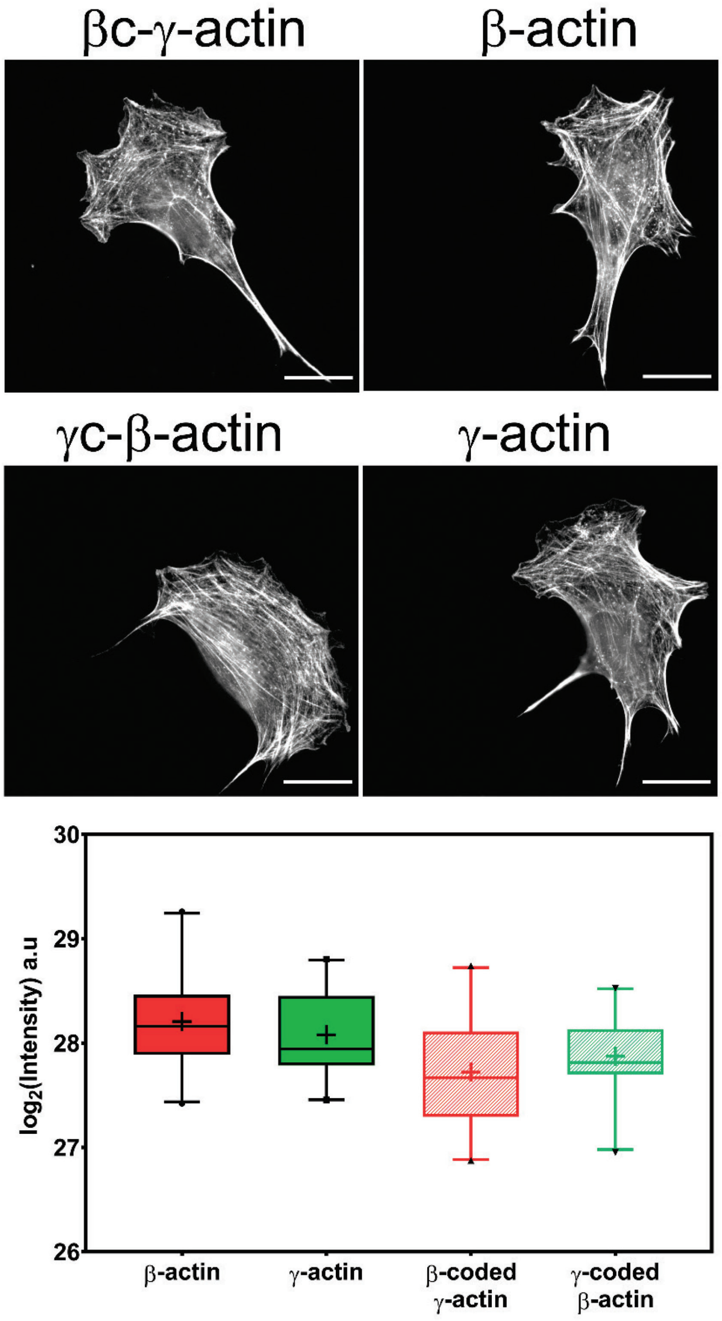


**Fig. S3, related to Fig.1. Cell transfection with actin isoforms does not perturb the level or distribution of the actin polymer.** A, Images of GFP-actin transfected cells lines visualized by fluorescent phalloidin staining. B, Box and whiskers plot of log2 (Phalloidin intensity) in each cell line with error bars indicating 95% CI, + indicating the mean. Scale bar = 20 μm. N = 20 each for β-actin, γ-actin, γ-coded β-actin and 22 for β-coded γ-actin.

**Vedula_et_al_Fig.S4**

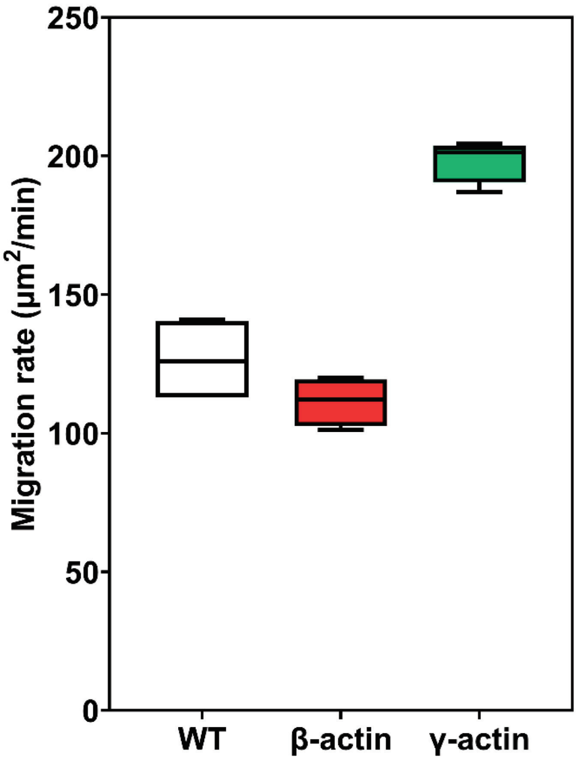


**Fig. S4, related to Fig. 1. Expressing γ-actin with β-actin 3′UTR increases cell migration rate.** Cell migration rate box plots showing untransfected (wildtype, WT), eGFP-β-actin (β-actin), and eGFP-γ-actin (γ-actin). N=4 for each cell line.

**Vedula_et_al_Fig.S5**

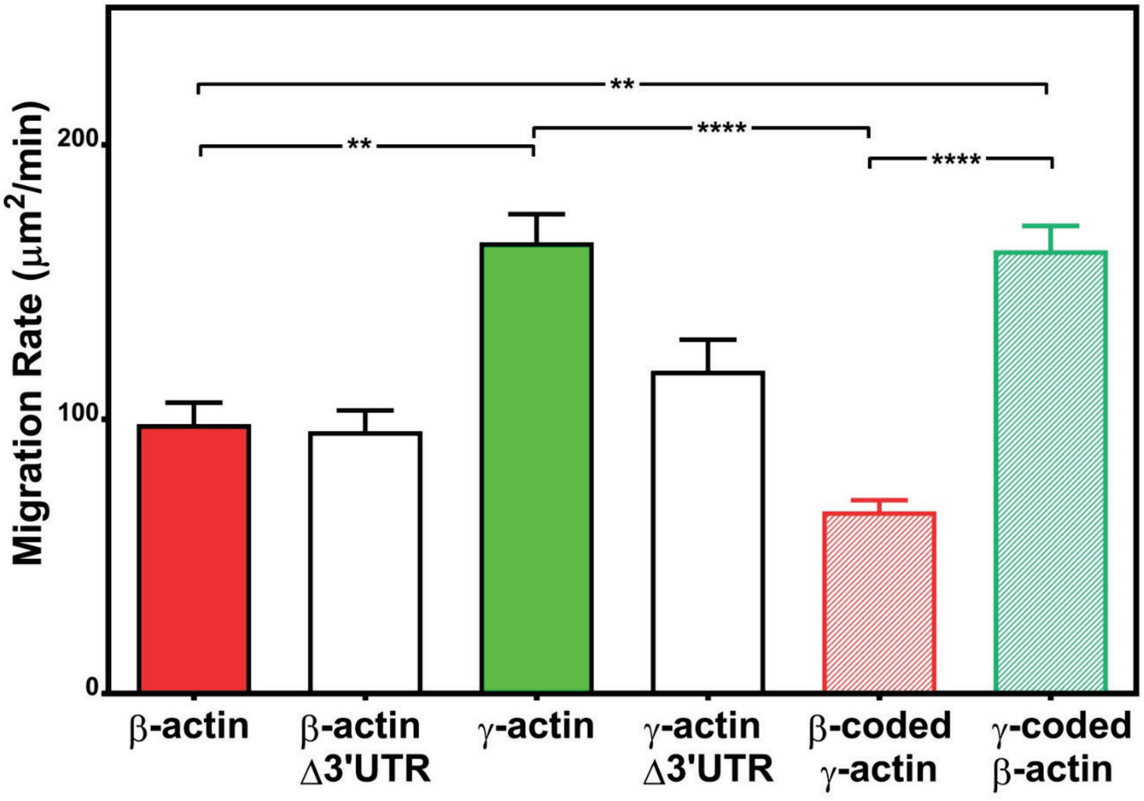


**Fig. S5, related to Fig. 1. Targeting of actin isoforms to the cell leading edge maximizes their effect on cell migration rates.** Rates plots of migration rates of cell lines expressing different actin constructs, derived as area covered over time by the entire cell monolayer as shown in Fig. 1. Δ3'UTR denotes the constructs lacking the β-actin 3'UTR responsible for zipcode targeting. All other constructs contained the β-actin 3'UTR. Cell migration rates were calculated as the area between the yellow and the green lines. Error rabs represent SEM. N=5 (for β-actin Δ3′UTR); 8 (for γ-actin Δ3′UTR); 20 (for β-actin); 22 (for γ-actin); 18 (for βc-γ-actin); 19 (for γc-β-actin). One way non-parametric ANOVA yielded a p value $<0.0001$ with multiple comparisons shown on the plot. $^{*}p<0.05$, $^{**}p<0.01$, $^{***}p<0.001$, and $^{****}p<0.0001$.

**Vedula_et_al_Fig.S6**

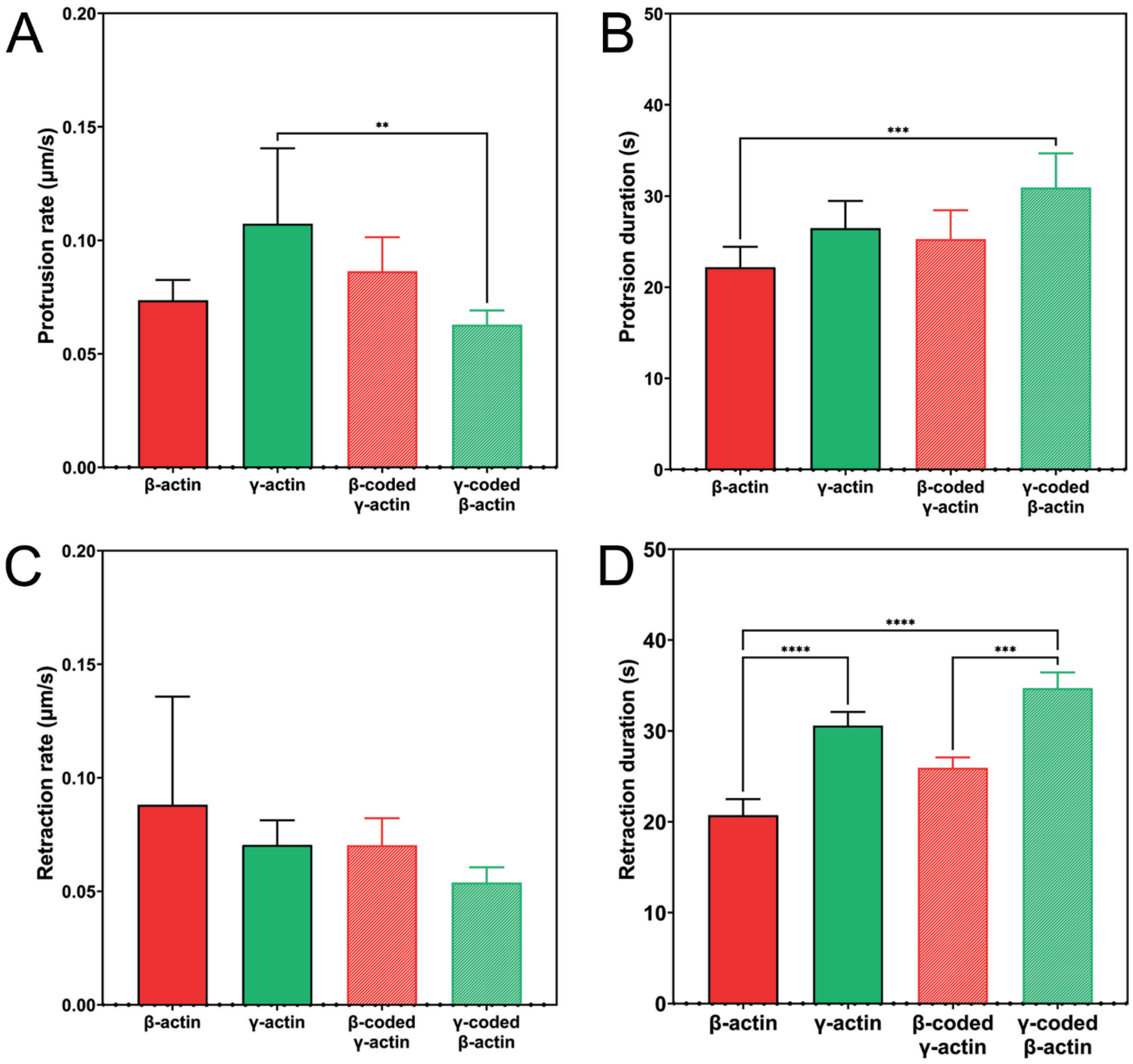


**Fig. S6, related to Fig. 2. Protrusion and retraction dynamics of cells at wound edge during wound healing.** A. Plot of protrusion rate measured from kymographs. B. Plot of protrusion duration. Number of protrusions analyzed: β-actin = 139, γ-actin = 130, β-coded-γ-actin = 143, γ-coded-β-actin = 121. C. Plot of retraction rate measured from kymographs. D. Plot of retraction duration. Number of retractions analyzed: β-actin = 112, γ-actin = 108, β-coded-γ-actin = 124, γ-coded-β-actin = 101. Error bars represent mean ± 95% CI. One-way parametric ANOVA yielded a p value = 0.0101 (protrusion rate), 0.0013 (protrusion duration), 0.3413 (retraction rate), and <0.0001 (retraction duration) with multiple comparisons shown on the plot. *p<0.05, **p<0.01, ***p<0.001, and ****p<0.0001.

**Vedula_et_al_Fig.S7**

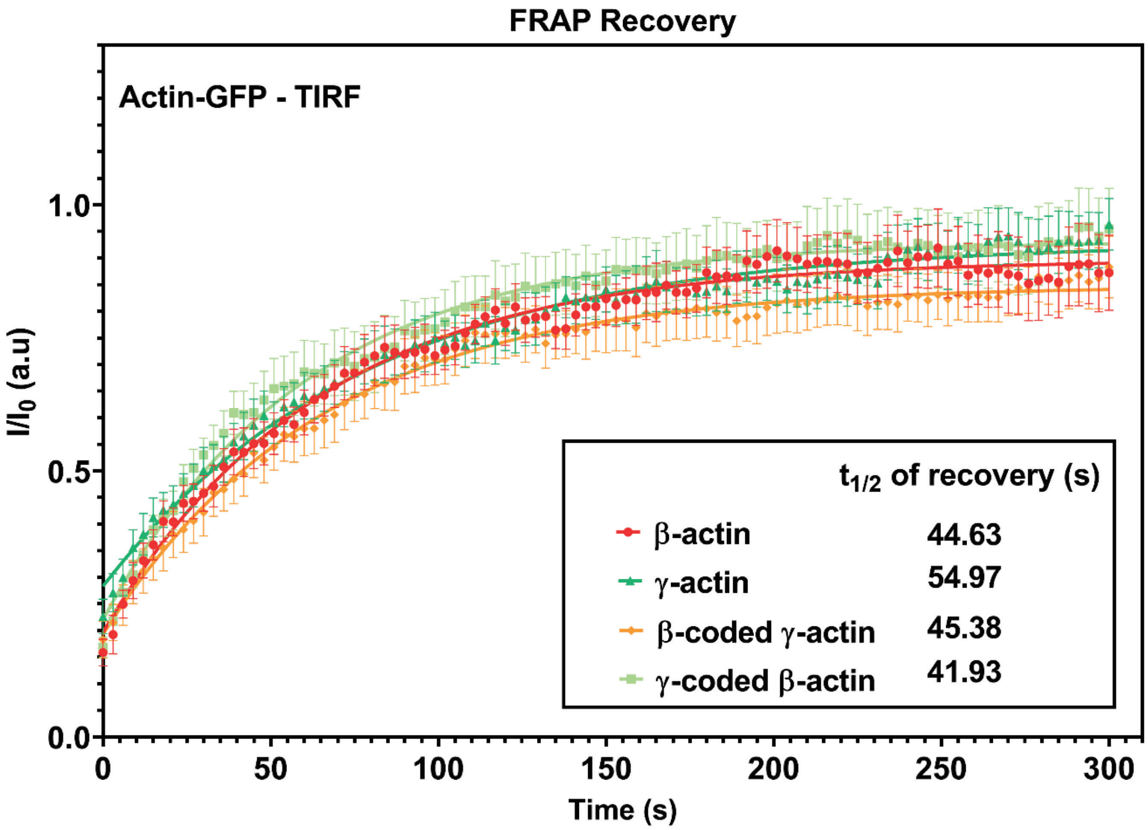


**Fig. S7, related to Fig. 2. FRAP recovery curves for actin patches in cells at wound edge during wound healing imaged in TIRF-M.** Fluorescence recovery after photobleaching (FRAP) curves for the recovery of actin bleached at the focal adhesions. $t_{1/2}$ of recovery for each actin variant is listed in the inset. N = 8 (β-actin), 10 (γ-actin), 10 (β-coded-γ-actin), and 9 (γ-coded-β-actin) actin patches. Each data set was fit to a one-phase association curve. R squared values ranged from 0.52 to 0.62. A single curve was unable to fit all data sets, indicating the recovery curves were different from each other (p-value < 0.0001). The half-life of recovery for each curve derived from the fit are β-actin = 44.63s, γ-actin = 54.97s, β-coded-γ-actin = 45.38s, and γ-coded-β-actin = 41.93s. Error bars are SEM.

**Vedula_et_al_Fig.S8**

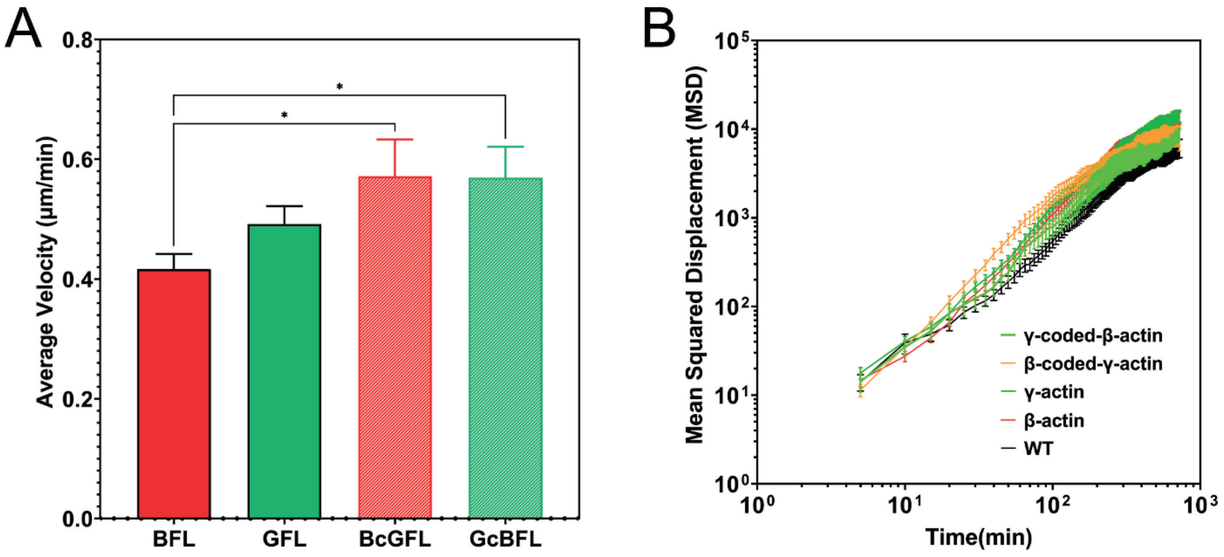


**Fig. S8, related to Fig.3. Single cell motility is not dependent on actin isoform coding or amino acid sequences.** A. Average velocity plots for single cell migration in sparse cultures over a 3 hour period. One-way parametric ANOVA yielded a p value = 0.0191 with multiple comparisons shown on the plot. *$p<0.05$. B. Log-log plots of Mean Squared Displacement vs time for each of the cell lines indicated on the graph. Number of cells: β-actin = 86, γ-actin = 121, β-coded-γ-actin = 97, γ-coded-β-actin = 69, untransfected (WT) = 100.

**Vedula_et_al_Fig.S9**

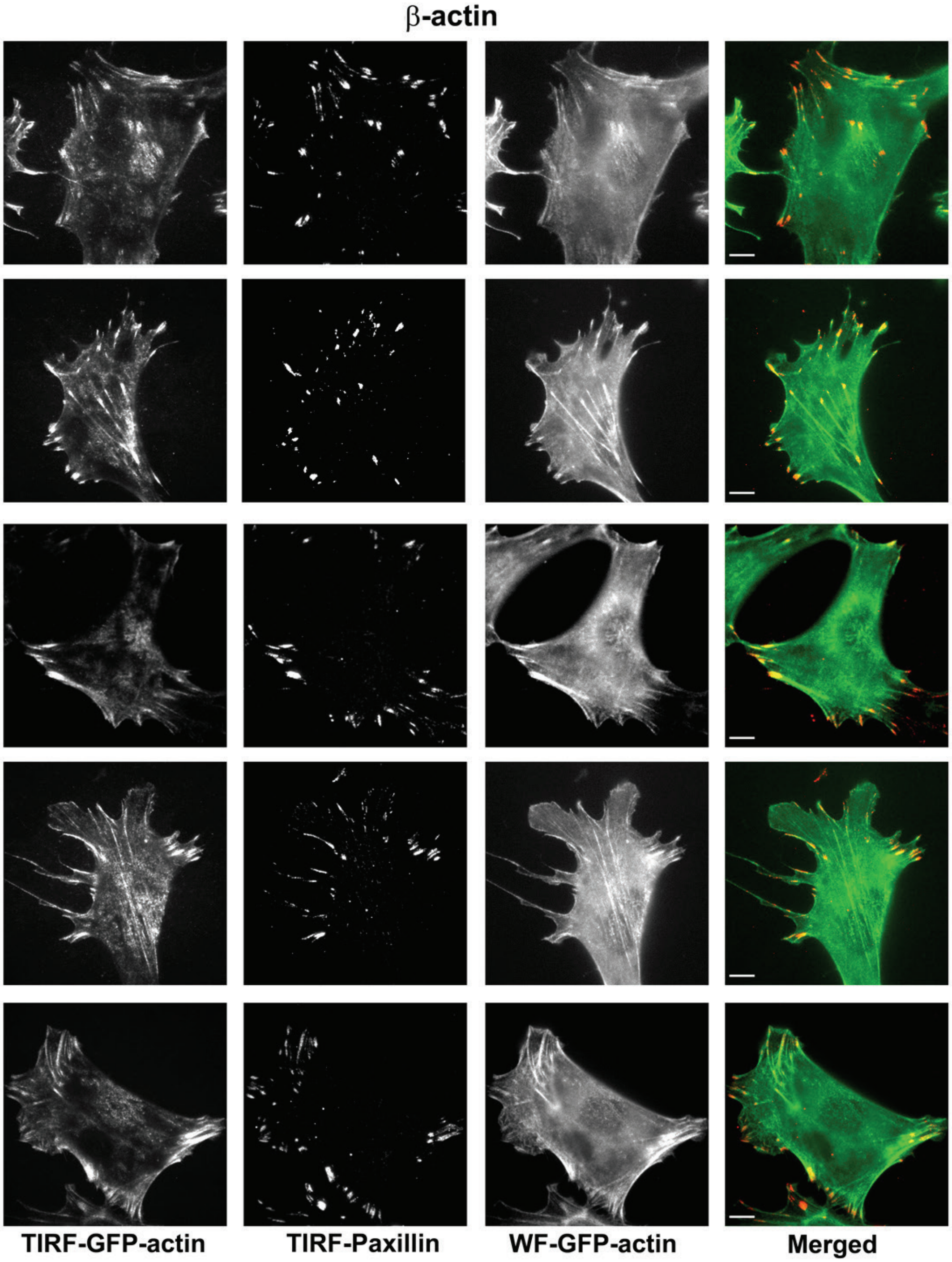


**Fig. S9, related to Fig.3. Representative images of paxillin and eGFP-actin distribution in cells stably transfected with eGFP-β-actin.** Scale bar, 10 μm.

**Vedula_et_al_Fig.S10**

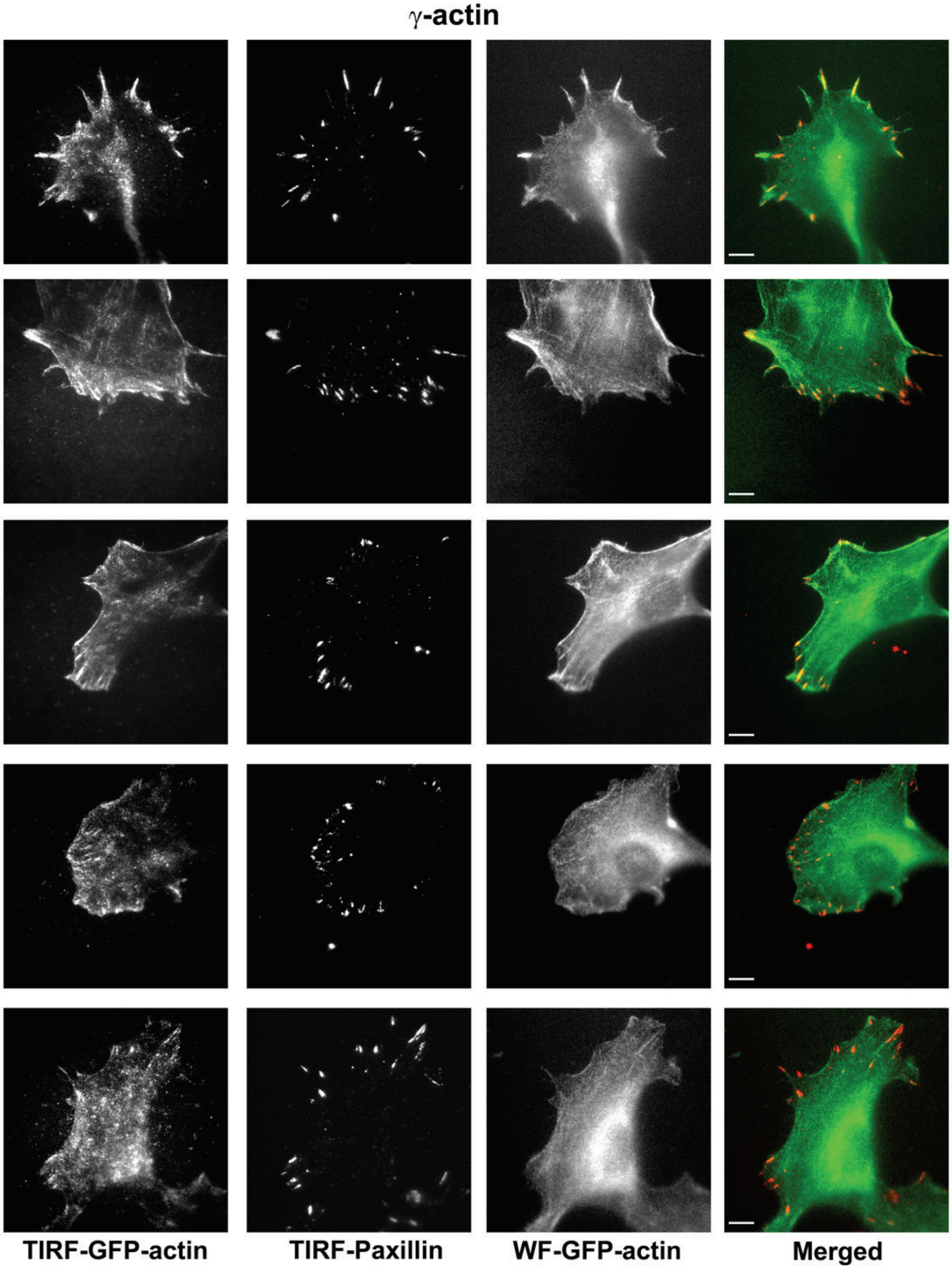


**Fig. S10, related to Fig.3. Representative images of paxillin and eGFP-actin distribution in cells stably transfected with eGFP-γ-actin.** Scale bar, 10 μm.

**Vedula_et_al_Fig.S11**

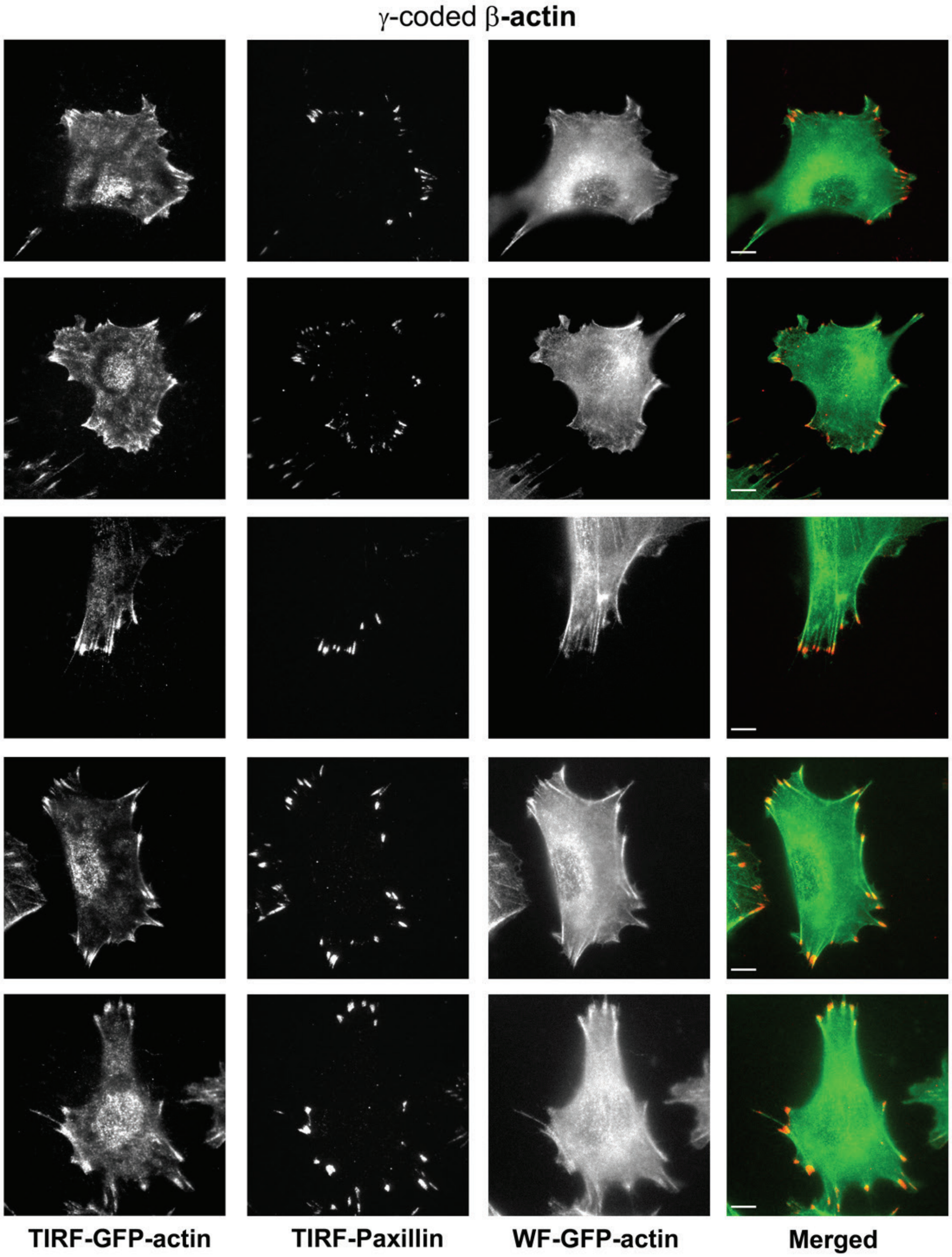


**Fig. S11, related to Fig.3. Representative images of paxillin and eGFP-actin distribution in cells stably transfected with eGFP-γ-coded β-actin.** Scale bar, 10 μm.

**Vedula_et_al_Fig.S12**

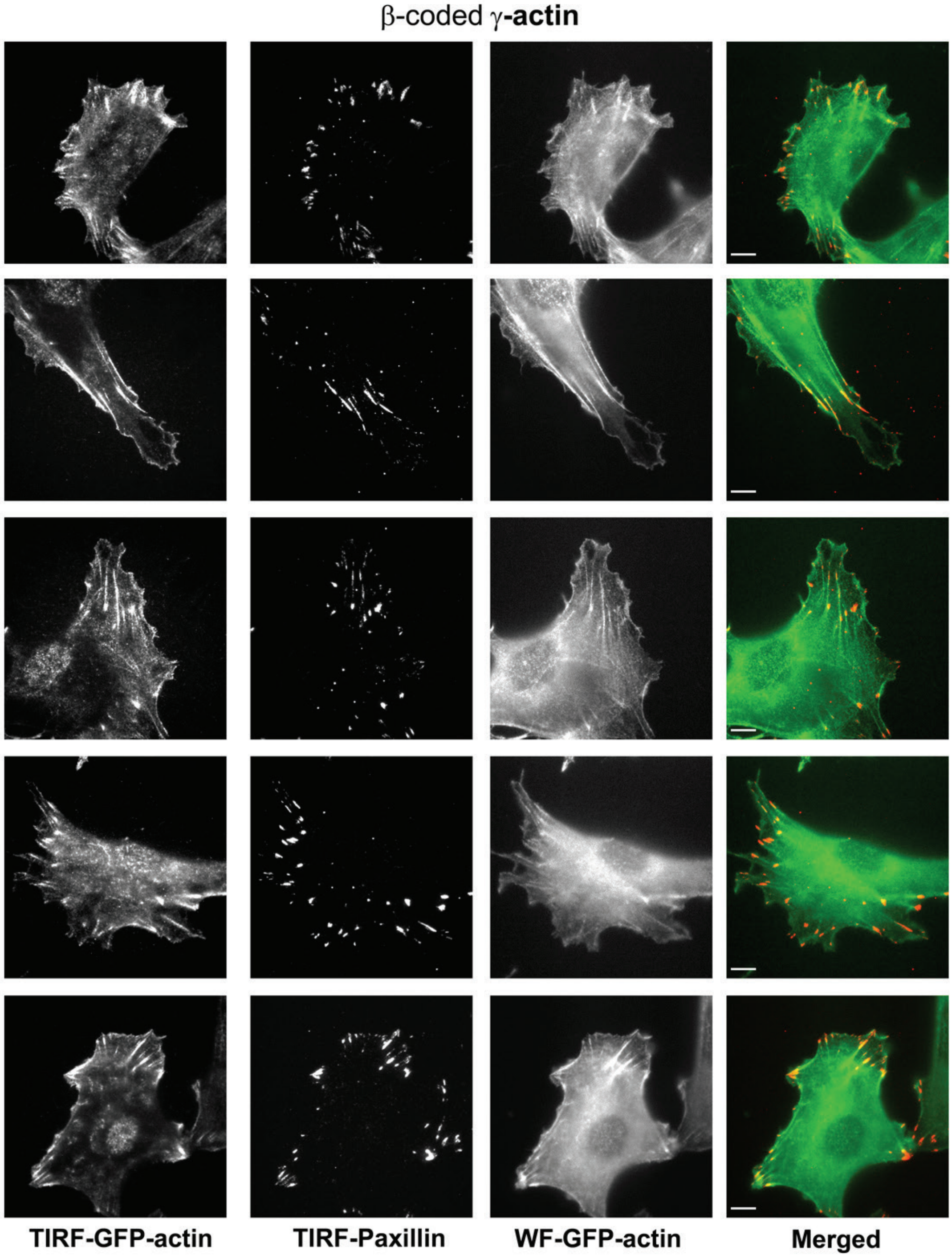


**Fig. S12, related to Fig.3. Representative images of paxillin and eGFP-actin distribution in cells stably transfected with eGFP-β-coded γ-actin.** Scale bar, 10 μm.

**Vedula_et_al_Fig.S13**

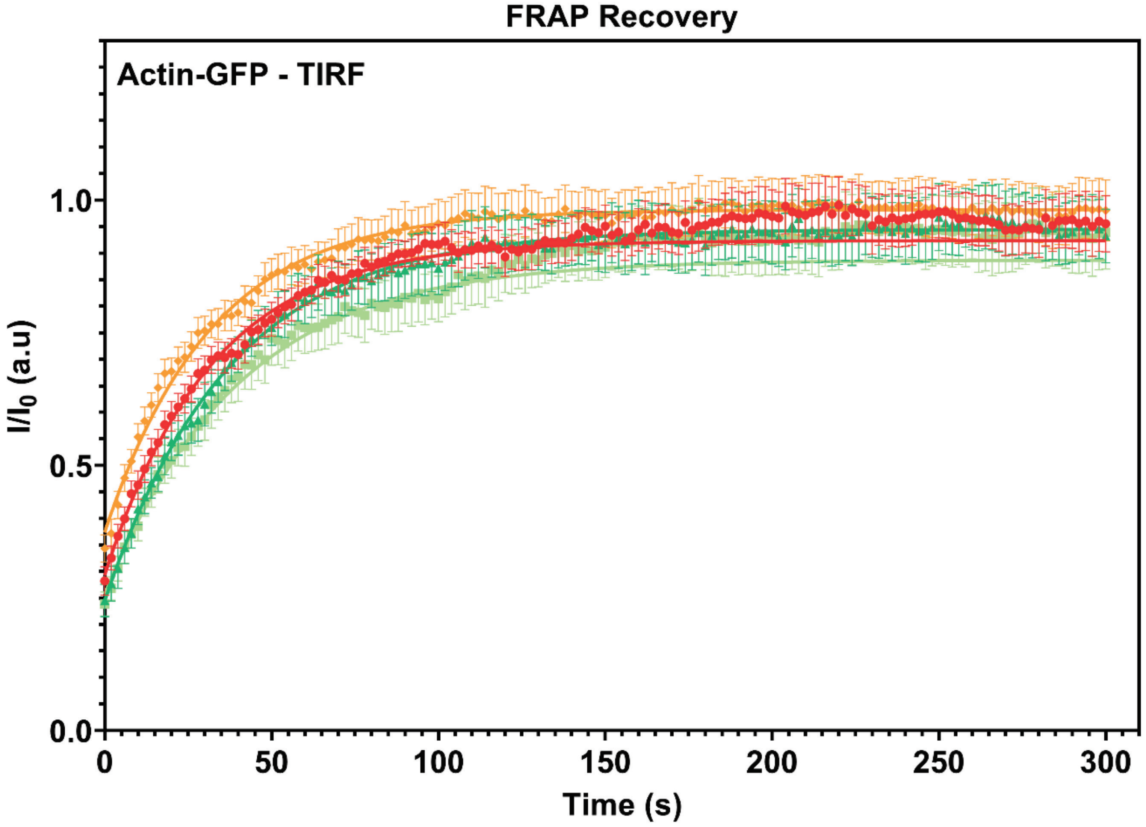


**Fig. S13, related to Fig.3. FRAP recovery curves for actin patches in single cell cultures imaged in TIRF-M.** Fluorescence recovery after photobleaching (FRAP) curves for the recovery of actin bleached at the focal adhesions. N = 15 actin patches for each cell line. Each data set was fit to a one-phase association curve. R squared values ranged from 0.36 to 0.59. A single curve was unable to fit all data sets, indicating the recovery curves were different from each other (p-value < 0.0001). The half-life of recovery for each curve derived from the fit are β-actin = 22.30s, γ-actin = 25.99s, β-coded-γ-actin = 22.13s, and γ-coded-β-actin = 27.87s. Error bars are SEM.

**Vedula_et_al_Fig.S14**

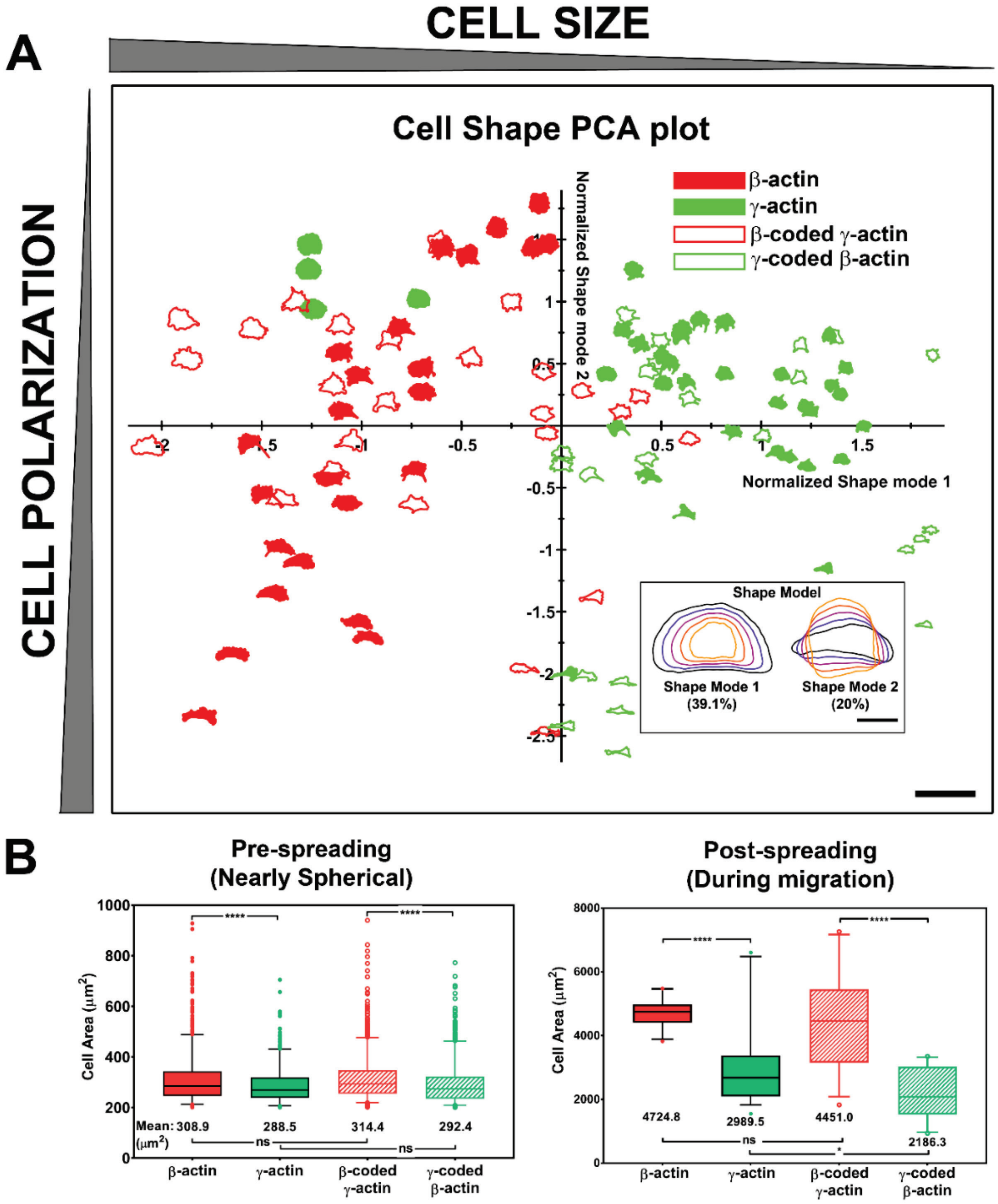


**Fig. S14, related to Fig.3 Actin isoforms confer differential effects on cell shape and spreading.** A, principle component analysis (PCA) plot classifying cell shapes from the four cell lines listed on the top right according to the shape modes shown in the insert on the bottom right. Shapes in the chart represent the actual cell outlines, with the size (Shape Mode 1) decreasing from left to right and the polarization variance (Shape Mode 2) spreading along the y axis. Scale bars, 250µm, and Inset, 50µm. B, quantification of the area or spherical trypsinized cells (left, N = 1230, 600, 1096, and 925) and the footprint of cell spread on the substrate (right, N = 25, 36, 27, and 23). Results for one-way ANOVA with multiple comparison are indicated on the graph.

**Vedula_et_al_Fig.S15**

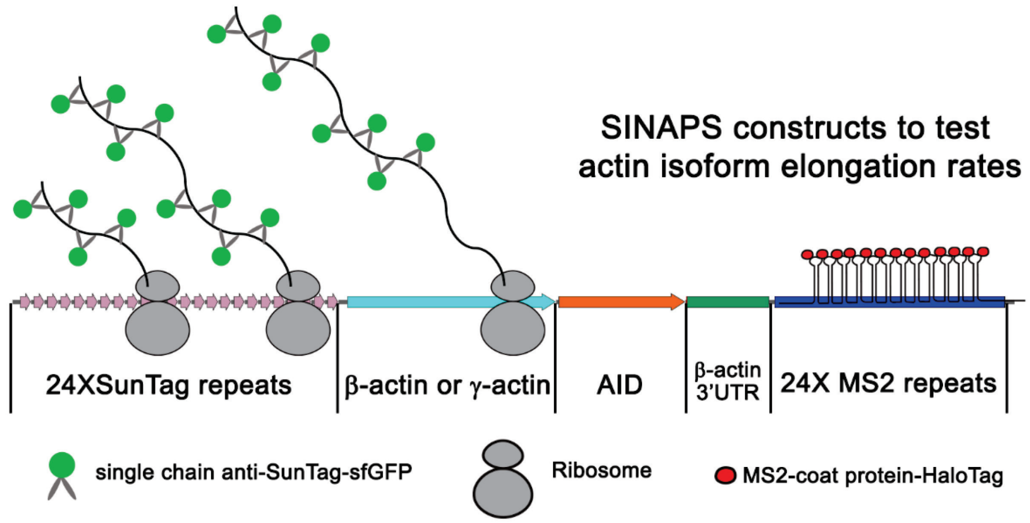


**Fig. S15, related to Fig.4B. Illustration of single molecule imaging of nascent peptide synthesis (SINAPs) for β- and γ-actin.** The 5′ end of the construct encodes 24 Suntag repeats, each of which upon translation bind to sfGFP-fused anti-Suntag antibody fragment expressed in the cells. This is followed by the β- or γ-actin coding sequence. This is then followed by Auxin induced degron, which once translated targets the nascent peptide for degradation. The construct also contains the β-actin 3′UTR required for mRNA zipcode mediated targeting to the leading edge. Lastly, the construct encodes 24 MS2 repeats each, which bind to MS2 coat binding protein fused to Halo-Tag. These MS2 repeats were used to visualize the mRNA of SINAPs constructs.

**Vedula_et_al_Fig.S16**

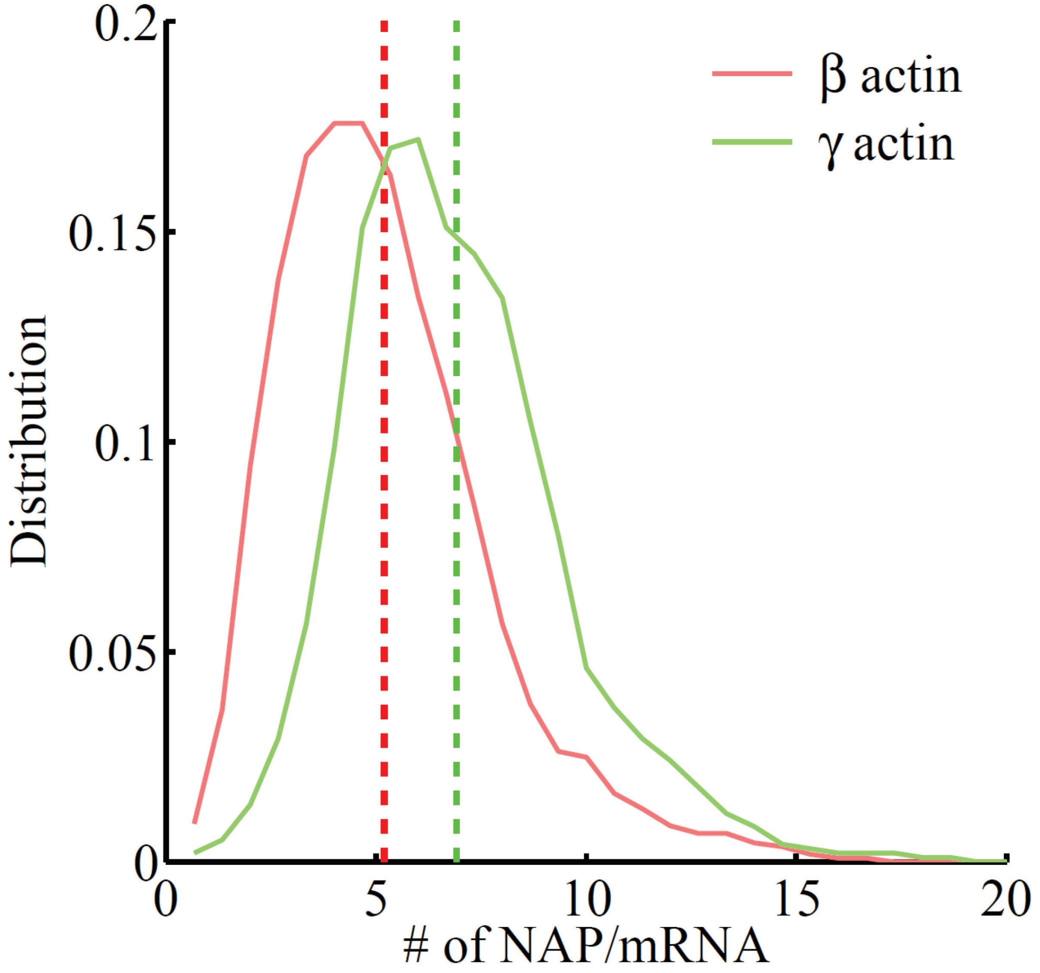


**Fig. S16, related to Fig. 4B.** Frequency distribution of number of nascent peptides (NAP) per mRNA on SINAPs constructs of β-actin (red) or γ-actin (green). Note the number of NAPs/mRNA is higher for γ-actin, indicating that γ-actin translation elongation is slower than that of β-actin.

**Vedula_et_al_Fig.S17**

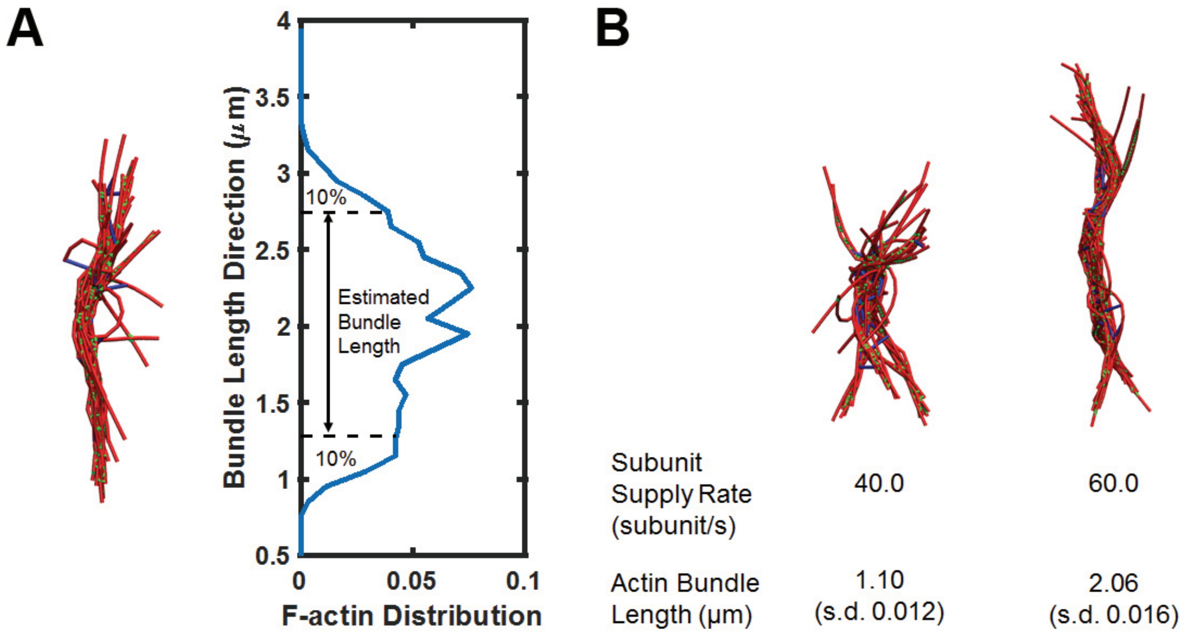


**Fig. S17, related to Fig.5. Simulations of actin bundle growth from the focal adhesion.** A. Measurements of the actin bundle length from distribution plots of F-actin distribution in the Z-axis. B. Representative simulation snapshots similar to those shown in Fig. 5A but with higher non-muscle myosin motor concentration (0.021 μM).

**Vedula_et_al_Table S1, related to Fig.5.**

| Parameter Name | Value | Reference and notes |
|---|---|---|
| Filaments | | |
| Filament polymerization and depolymerization rates | $k_{on} = 12.9\ (11.6 + 1.3) \mu M^{-1} s^{-1}$<br>$k_{off} = 2.2 (1.4 + 0.8)\ s^{-1}$ | (Fujiwara et al., 2007) Taking both barbed end and pointed ends into account |
| Filament bending constant | $672.5\ pN \cdot nm$ | (Ott et al., 1993) Bending constant between connecting cylinders* |
| Filament stretching constant | $100.0\ pN/nm$ | (Popov et al., 2016) Stretching constant of cylinder |
| Motors | | |
| Binding rate constant | $0.2\ s^{-1}$ per head | (Stam et al., 2015) |
| Duty ratio | 0.1 | (Stam et al., 2015) |
| Number of motor heads per mini-filament | 15-30 | (Verkhovsky and Borisy, 1993) |
| Stretching constant | $56.0\ pN/nm$ | (Popov et al., 2016) |
| Characteristic stall force | $15.0\ pN$ per head | (Popov et al., 2016) |
| Characteristic motor unbinding force | $12.6\ pN$ per head | (Erdmann et al., 2013) |
| Crosslinkers | | |
| Binding rate constant | $0.7\ \mu M^{-1} s^{-1}$ | (Wachsstock et al., 1993) |
| Unbinding rate constant | $0.3\ s^{-1}$ | (Wachsstock et al., 1993) |
| Stretching constant | $8.0\ pN/nm$ | (DiDonna and Levine, 2007) |
| Characteristic linker unbinding force | $17.2\ pN$ | (Ferrer et al., 2008) |

* Filaments are discretized into cylinders, and the length cylinder ranges from 2.7 nm (1 subunit) to a maximum 108 nm (40 subunits). Bending is only allowed between two connecting cylinders.

**Supplemental Video Legends:**

Video 1: Migration of β-actin-transfected cells.

Video 2: Migration of γ-actin-transfected cells.

Video 3: Migration of β-coded γ-actin-transfected cells.

Video 4: Migration of γ-coded β-actin-transfected cells.

Video 5: Simulation of actin bundle growth from a focal adhesion at a slower subunit supply rate.

Video 6: Simulation of actin bundle growth from a focal adhesion at a faster subunit supply rate
